# Tesofensine, a novel antiobesity drug, silences GABAergic hypothalamic neurons

**DOI:** 10.1101/2023.08.02.551706

**Authors:** Claudia I. Perez, Jorge Luis-Islas, Axel Lopez, Xarenny Diaz, Omar Molina, Benjamin Arroyo, Mario G. Moreno, Elvi Gil Lievana, Esmeralda Fonseca, Gilberto Castañeda-Hernández, Ranier Gutierrez

## Abstract

Obesity is a major global health epidemic that has adverse effects on both the people affected as well as the cost to society. Several anti-obesity drugs that target GLP-1 receptors have recently come to the market. Here we describe the effects of tesofensine, a novel anti-obesity drug that acts as a triple monoamine neurotransmitter reuptake inhibitor. We investigated its effects on weight loss and underlying neuronal mechanisms in mice and rats using various techniques. These include behavioral tasks, DeepLabCut videotaped analysis, electrophysiological ensemble recordings, optogenetic activation, and chemogenetic silencing of GABAergic neurons in the Lateral Hypothalamus (LH). We found that tesofensine induced greater weight loss in obese than lean rats, which was associated with changes in LH ensemble activity. In Vgat-ChR2 and Vgat-IRES-cre transgenic mice, we found for the first time that tesofensine inhibited a subset of LH GABAergic neurons, reducing their ability to promote feeding behavior, and chemogenetically silencing them enhanced tesofensine’s food-suppressing effects. Unlike phentermine, a dopaminergic appetite suppressant, tesofensine causes few, if any, head-weaving stereotypy at therapeutic doses. Most importantly, we found that tesofensine prolonged the weight loss induced by 5-HTP, a serotonin precursor, and blocked the body weight rebound that often occurs after weight loss. Behavioral studies on rats with the tastant sucrose indicated that tesofensine’s appetite suppressant effects are independent of taste aversion and do not directly affect the perception of sweetness or palatability of sucrose. In summary, our data provide new insights into the effects of tesofensine on weight loss and the underlying neuronal mechanisms, suggesting that tesofensine may be an effective treatment for obesity and that it may be a valuable adjunct to other appetite suppressants to prevent body weight rebound.

## INTRODUCTION

Obesity is a worldwide health problem that has reached epidemic proportions. Although diet and exercise are the primary treatments for obesity, these activities are often supplemented using appetite suppressants. Tesofensine (NS2330) is a triple monoamine re-uptake inhibitor with an affinity for dopamine (DAT), serotonin (SERT), and norepinephrine (NET) transporters. It has been shown to have antiobesity effects in animal and human studies (Astrup et al., 2008a). Tesofensine significantly reduced daily food intake in rats under a 16-day treatment regimen, leading to a significant and sustained decrease in body weight. However, the anorexigenic effect of tesofensine progressed to tolerance, while the weight loss effect did not (Axel et al., 2010). Hence, tesofensine is a dual-action drug with anorexigenic and metabolic properties, increasing energy expenditure. More impressively, tesofensine is more effective at reducing body weight in high-fat-fed rats than in chow-fed rats (Axel et al., 2010; Hansen et al., 2013). Moreover, it is known that tesofensine activates α1 adrenergic receptors and, to a lesser extent, dopamine D1 receptors (Appel et al., 2014; Axel et al., 2010; Hansen et al., 2013). exhibits potent antiobesity effects, but the underlying cellular mechanisms are not yet fully elucidated. This study first aims to identify the neuronal correlates of tesofensine-induced weight loss in the Lateral Hypothalamus (LH) in lean and obese rats.

The LH is a brain region that regulates numerous physiological processes involving seeking and feeding behaviors (Panksepp, 2004). Lesions in the LH can cause decreased food intake and weight loss, while stimulation can increase food intake and promote obesity (Anand and Brobeck, 1951; Hoebel and Teitelbaum, 1962). The LH comprises two major neuronal populations GABAergic and glutamatergic neurons, that play opposing and bidirectional roles in reward and feeding (Jennings et al., 2015; Rossi, 2023; Rossi et al., 2019). In mice and primates, activation of LH GABA neurons promotes food intake, while silencing them inhibits food intake (Garcia et al., 2021; Ha et al., 2023; Nieh et al., 2015). In contrast, in mice, the activation of LH glutamatergic neurons inhibits food intake, while their inhibition promotes food intake (Rossi et al., 2019). However, it is currently unknown whether tesofensine targets these neuronal populations.

A second aim of this study, in mice, is to characterize how tesofensine targets LH GABAergic neurons to modulate feeding behavior. A third aim was to compare in lean rats the anti-obesity effects of tesofensine with phentermine, another appetite suppressant that increases dopamine efflux in the nucleus accumbens and also induces head weaving stereotypy (Baumann et al., 2000; Kalyanasundar et al., 2015). We also investigated the pharmacological interaction between tesofensine and 5-HTP, a serotonin precursor and appetite suppressant, and found that tesofensine delayed weight loss rebound (Amer et al., 2004; Birdsall, 1998; Halford et al., 2007). Finally, we investigated whether tesofensine affects the gustatory perception of sweetness, as it is reported to decrease craving for sweet food (Gilbert et al., 2012). Overall, our study provides insights into the potential use of tesofensine as an effective treatment for obesity.

## RESULTS

### Tesofensine demonstrated greater weight loss efficacy in obese rats

First, we asked whether tesofensine has the same efficacy in obese and lean rats. To do this, we characterized the anorexigenic and body weight loss effects induced by tesofensine in both types of rats. Rats had ad libitum access to a standard chow or a high-fat diet for 12 weeks during childhood and adolescence. Obese rats fed with a high-fat diet weighed more initially (588.7 ± 14.1 g) than rats on a standard chow diet (497.8 ± 15.1 g, *p* < 0.05) (**Figure 1A**, see inset). Every 24 hours, we weighed the daily food intake and body weight and subsequently injected tesofensine or saline. A constant weight gain was observed in Chow-Saline and HFD-Saline groups (**Figure 1A**). In contrast, after three days of treatment, Chow-Tesofensine and HFD-Tesofensine groups decreased body weight compared to control rats receiving saline. Afterward, the HFD-Tesofensine group lost weight faster than Chow-tesofensine-treated rats. Lean rats treated with tesofensine reached a plateau after the third day until the end of treatment and demonstrated less body weight loss than obese animals treated with tesofensine. This study found that tesofensine was more effective in inducing weight loss in obese rats than in lean rats.

**Figure 1.**
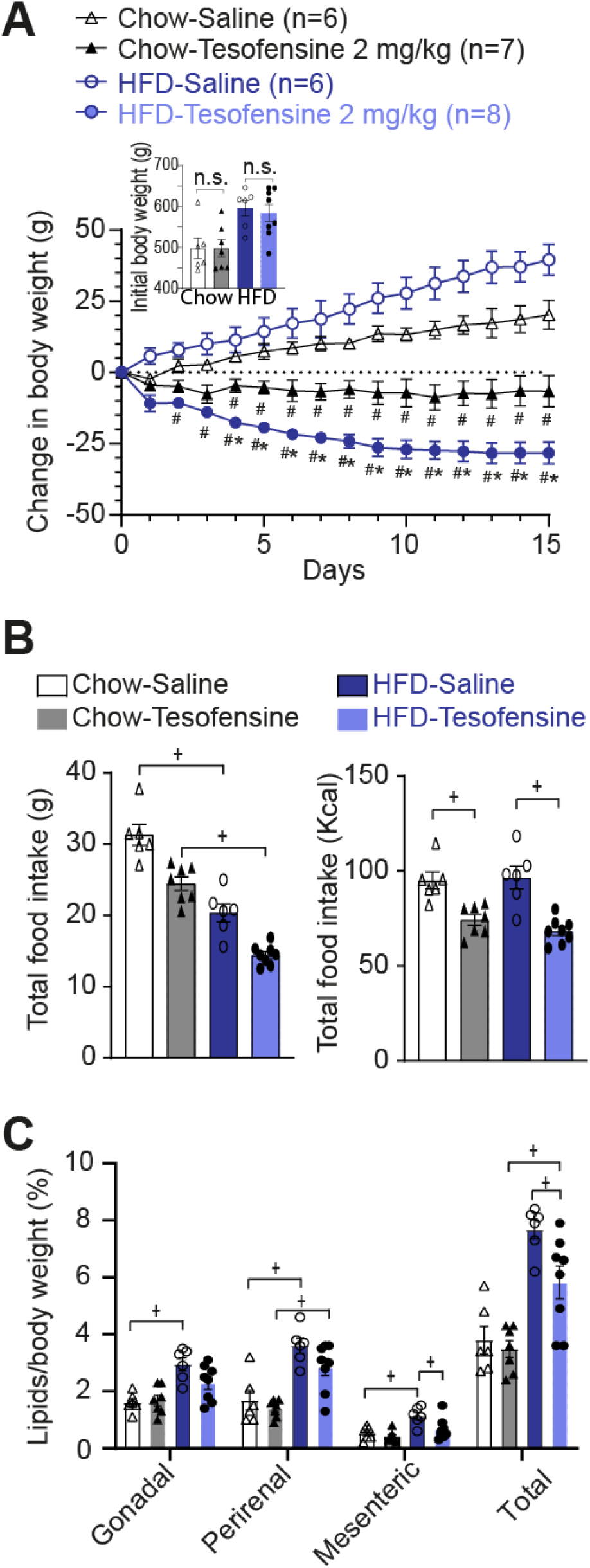
Tesofensine induces greater weight loss in HFD-fed rats compared to chow-fed rats. **A**. Change in body weight across treatment days with the treatment groups receiving subcutaneous injections of tesofensine (2 mg/kg) and the control groups receiving saline. The inset graph shows the body weight of chow-fed and HFD-fed rats before treatment. No significant difference on body weight was present on both groups fed with chow or between both groups fed with HFD. **B.** Total grams of food consumed per group (left panel) and total caloric intake (right panel). Both groups treated with tesofensine consumed fewer calories than saline control groups. **C.** Visceral fat content (%) relative to body weight in gonadal, perirenal, and mesenteric deposits. Data are presented as mean ± SEM. The symbols #, *, and + indicate statistical significance at *p* < 0.05 compared with Chow-Saline, Chow-Tesofensine, and significantly different between groups, respectively.

The weighed food intake is shown in **Figure 1B**, left panel. Since diets have different amounts of energy, we normalized them to kilocalories (**Figure 1B**, right panel). It can now be seen that tesofensine reduced the total energy intake in both Chow-Tesofensine and HFD-Tesofensine treated groups compared with control rats receiving saline (**Figure 1B**, right panel). We observed no significant difference in kilocalories consumed between Chow-Tesofensine vs. HFD-Tesofensine groups (*p* = 0.6). The results suggest that the increased effectiveness of tesofensine in promoting weight loss in obese rats could be because of an increase in energy expenditure rather than its anorexigenic effects.

Next, we quantified the effect of tesofensine on the visceral fat proportion of body weight in lean and obese rats. We found a significant difference in total visceral fat (composed of gonadal, perirenal, and mesenteric fat) between the HFD-Saline and HFD-Tesofensine groups (**Figure 1C**). However, the total fat in the Chow-Tesofensine group did not differ significantly from that of the Chow-Saline group. These results indicate that tesofensine reduced total visceral fat, mainly mesenteric fat deposits, in obese rats.

### Tesofensine-induced modulation of lateral hypothalamic neurons is more pronounced in obese than in lean rats

The LH plays a vital role in seeking food and regulating feeding behavior (Garcia et al., 2021; Nieh et al., 2015; Panksepp, 2004). It is believed to be a primary target for various appetite suppressants, and recently it was found that tesofensine could be a potential treatment for hypothalamic obesity, a rare feeding disorder (Astrup et al., 2008a; Huynh et al., 2022; Saniona, 2022). Hence, LH might be a potential target of this drug.

To examine this possibility, we evaluated whether tesofensine exerts a differential effect on LH cell activity of lean versus obese subjects. Thus, we conducted multichannel recordings in the LH of six rats, three fed with a standard diet and three high-fat diet. During the recordings in the LH, we administered saline and tesofensine subcutaneously via a catheter. A total of 343 neurons in the Chow-fed rats and 361 in HFD-fed rats were recorded (**Figure 2A**). We then used a t-distributed stochastic neighbor embedding (t-SNE) to classify the firing rates of recorded neurons into ensembles. The results are shown in **Figure 2 Supplementary 1**, with the left panel depicting neurons recorded in Chow-fed rats and the right panel for neurons recorded in HFD-fed rats. **Figure 2B** shows the neuronal responses over two hours of recordings. The normalized firing rates were then color-coded, with blue indicating decreased and red increased activity. t-SNE analysis unveiled four ensembles: The first ensemble (E1) includes a cluster of neurons that exhibited a robust inhibition lasting 1.5 hours after the tesofensine administration. The second neuronal ensemble (E2) showed modest inhibition in activity. In contrast, the third and fourth ensembles showed modest (E3) and substantial (E4) neuronal responses (activation), respectively. The proportion of neurons in each ensemble differed significantly between obese and lean rats. Specifically, the proportion of neurons inhibited by tesofensine (E1) increased from 25% (86/343) in Chow-fed rats to 35% (125/361) in HFD-fed rats (Chi-square=7.19, *p*=0.0073). Similarly, tesofensine recruited a greater proportion of activated neurons (E4) in obese rats (31%, 113/361) than in lean rats (11%, 38/343) (Chi-square = 41.50, *p*<1.1771e-10) (**Figure 2C**). These findings indicate that administering tesofensine to rats exposed to a high-fat diet during childhood and adolescence resulted in a different modulation pattern in the LH compared to age-matched lean rats. In addition, the recorded population activity of neurons demonstrated a bias towards activation in obese rats but not in lean rats (**Figure 2D**). For example, the sum of ensembles 3 and 4, which exhibited increased firing rates after the tesofensine administration, was 56.2% (E3=90 + E4=113; 203/361) HFD-fed rats, compared to 46% (E3=120 + E4=38; 158/343) in Chow-fed rats (Chi-square=6.87, *p*=0.0087). Our analysis found that the differences between the effects of tesofensine on the LH ensemble activity of chow-fed and HFD-fed rats were statistically significant. These findings indicate that tesofensine effect over LH modulation is different in obese rats chronically exposed to a high-fat diet during childhood and adolescence, compared to lean subjects: tesofensine promotes the recruitment of cells belonging to strong-inhibitory (E1) and -excitatory (E4) responsive ensembles, and an increased sustained overall LH population activity.

**Figure 2.**
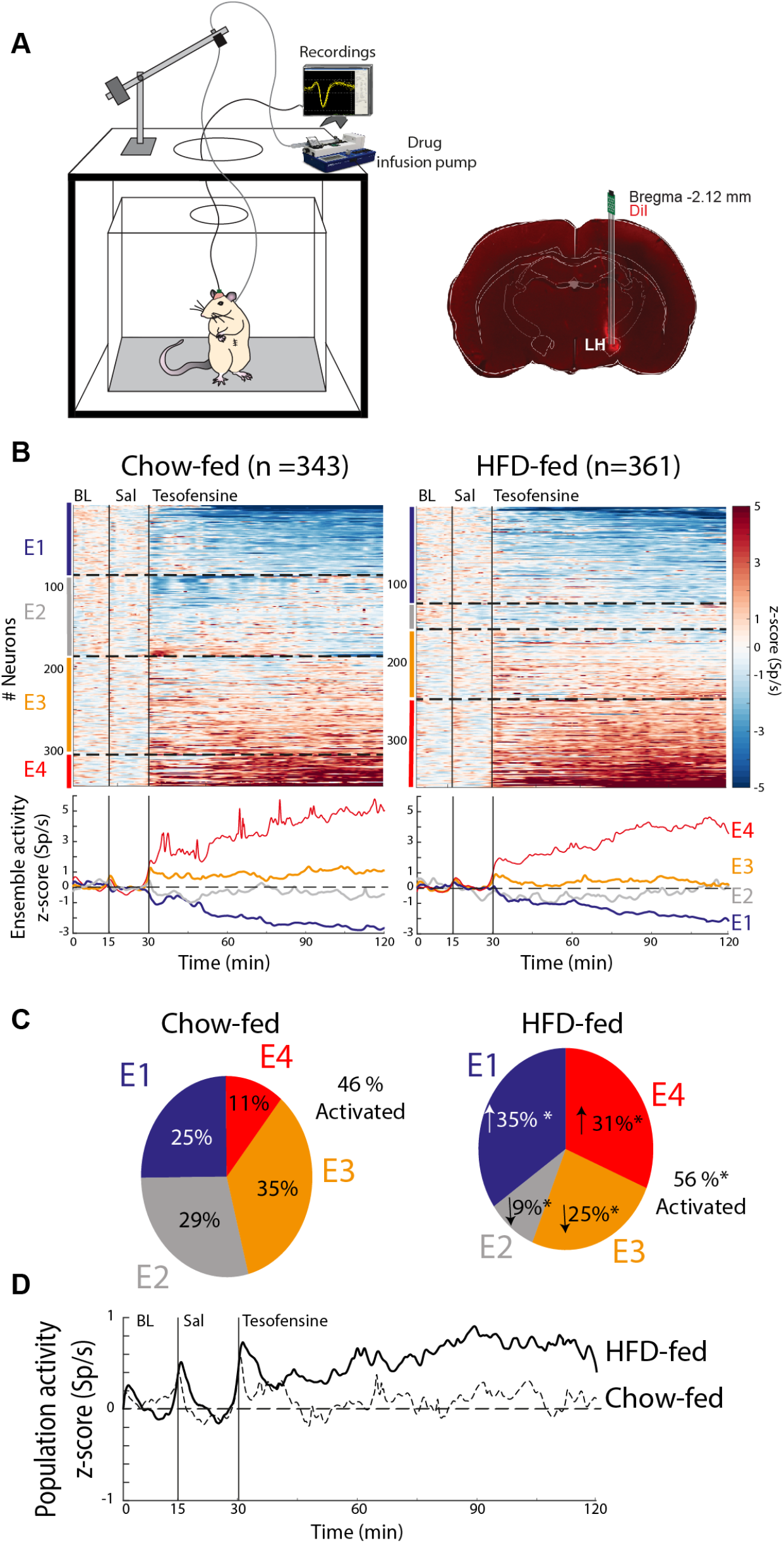
Tesofensine differentially modulates lateral hypothalamic neurons in lean and obese rats. **A.** Left: A schematic representation for extracellular recording in a freely moving rat and pump for automatic drug delivery. No food was available during recordings. Right: histology, identifying recording site in LH using a coronal section. Dil, a fluorescent and lipophilic dye, is used to mark the tip of the electrode track. **B.** t-SNE analysis of firing rates was used to group similar activity patterns into ensembles (**Figure 2, Supplementary fig. 1**). All neurons recorded were categorized into four ensembles (E1-E4) for Chow-fed rats (left) and HFD-fed rats (right). Normalized color-coded activity of each neuron over time is presented for chow-fed and HFD-fed rats, with red and blue indicating higher and lower z-score activity, respectively. Black vertical lines show the baseline (BL from 0-15 minutes), saline (Sal 15-30 minutes), and tesofensine administered at 30 minutes and recordings that lasted up to 120 minutes. Peri-Stimulus Time Histograms (PSTHs) below demonstrate the average neuronal ensemble activity, with dashed lines dividing each cluster (ensemble). **C.** Pie charts depict the percentage of neurons in each ensemble for Chow-fed and HFD-fed rats. Chi-square analysis showed a significant difference compared to the same ensemble in Chow-fed rats (* p<0.05). **D.** The z-score normalized population activity of all lateral hypothalamic neurons recorded in HFD-fed (n=361) and chow-fed rats (n=343) is presented.

### Tesofensine silenced LH GABAergic neurons in transgenic mice

Following the observation of distinct effects of tesofensine on LH activity in obese and lean rats, we investigated the specific cell type in this region that was primarily affected by the drug in mice. We hypothesize that tesofensine could affect GABAergic neurons due to its role in seeking and consummatory behaviors (Garcia et al., 2021; Nieh et al., 2015). To optogenetically identify LH-GABAergic neurons, we perform optrode recordings in lean Vgat-IRES-Cre mice, as depicted in **Figure 3A**. We recorded LH multichannel activity during a baseline period of at least 5 minutes before injecting saline or tesofensine 2 mg/kg subcutaneously on alternating days. After a minimum of 30 minutes, we conducted an optotagging assay comprising 5-minute blocks of active (50 Hz and laser turned 2s on, 4s off) and inactive periods. **Figure 3B** shows the firing rates of two distinct LH neurons. The first neuron exhibited a gradual decrease in firing rate following tesofensine administration. During the optotagging epoch, we identified it as GABAergic because it showed increased activity during the 5-minute block of photostimulation. Conversely, the second example is a non-GABAergic neuron because it was inhibited during photostimulation. Additionally, it exhibited a significant increase in firing rates following tesofensine administration. **Figure 3C** shows the color-coded activity of all neurons opto-identified as GABAergic and non-GABAergic and their population activity. During saline injection days (left panel), neither GABAergic nor non-GABAergic neurons were modulated after saline injection. During optotagging (see 33-66 minutes), only GABAergic neurons (blue trace) responded during laser stimulation.

**Figure 3.**
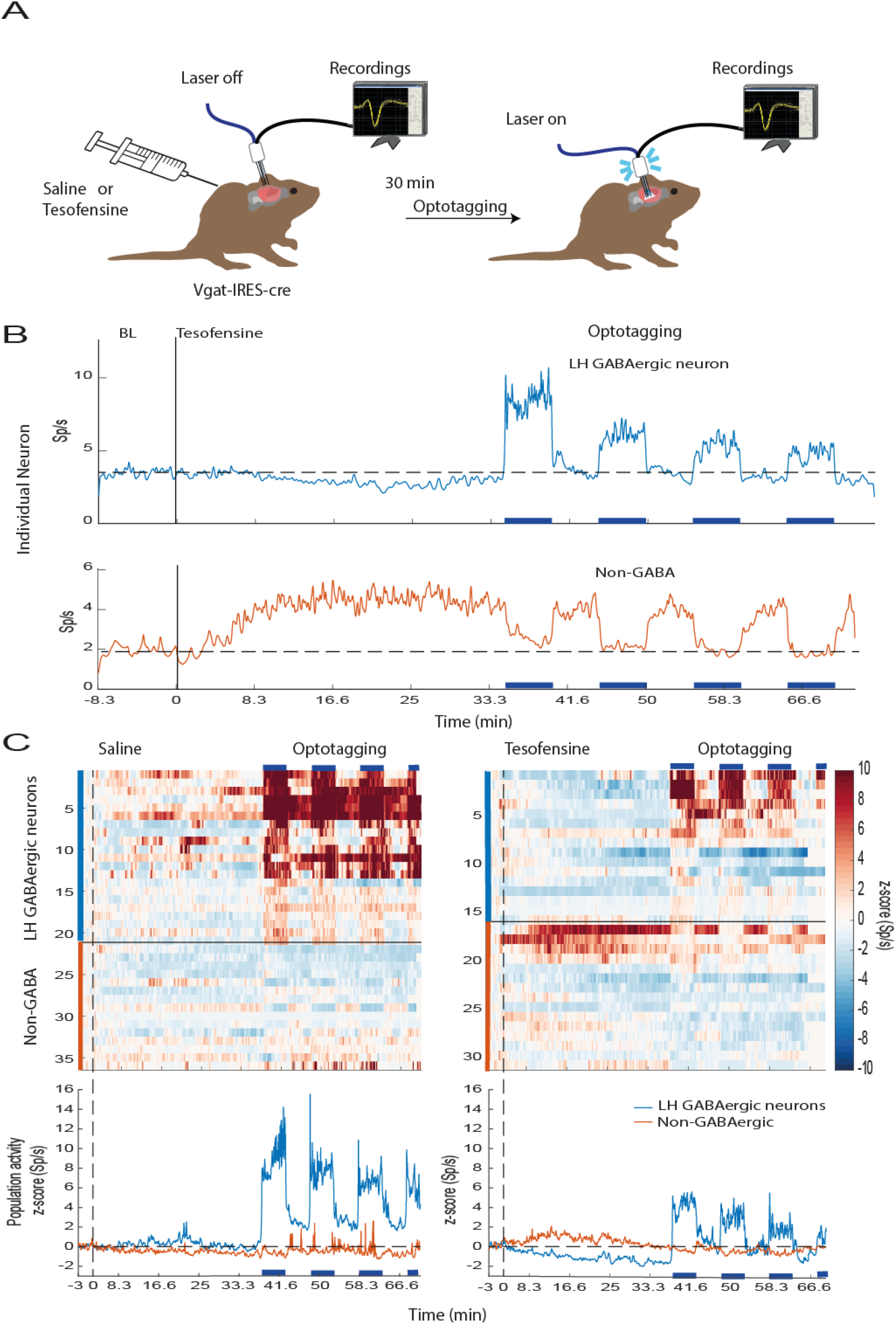
Tesofensine silences mice LH GABAergic neurons and attenuates their optogenetic activation. **A.** Extracellular neuronal activity was recorded in mice over a baseline period of 5 minutes before they received subcutaneous injections of saline or tesofensine (2mg/kg). After 30 minutes, a laser was activated on an open loop protocol. It was started with 5-minute blocks of active (50 Hz; 2s on + 4s off) and inactive periods (no laser). **B.** Upper, it depicts responses of single GABAergic neuron inhibited after tesofensine administration. Bottom, a non-GABAergic neuron. **C.** A heat map displaying neuronal activity during saline (left) and tesofensine (right) conditions. Blue horizontal lines indicate stimulation blocks. Below, the corresponding peri-stimulus time histogram (PSTH) shows the average activity of neurons during the stimulation blocks with the dashed line representing a zero z-score. Tesofensine reduced neuronal activity even during optogenetic stimulation, highlighting its ability to silence GABAergic neurons in the LH.

During sessions with tesofensine administration (right panel), we observed that the population activity of GABAergic neurons was gradually inhibited after tesofensine administration (blue trace, see 0-32 minutes). Moreover, the population responses elicited by the optogenetic stimulation (optotagging see 33-66 minutes) were significantly weaker under tesofensine treatment than saline injection (compare the blue PSTH lines in the right vs. left panels, respectively). In contrast, the non-GABAergic neurons (red trace) showed a slight increase in activity after tesofensine administration (see 0-32 mins). They were inhibited or not modulated during the optotagging assay (see 33-66 mins). Thus, tesofensine silences a subset of LH GABAergic neurons and attenuates their open loop optogenetic activation.

### Tesofensine reduced feeding behavior induced by optogenetic activation of LH GABAergic neurons in lean Vgat-ChR2 mice

Based on the findings that tesofensine attenuates LH-GABAergic activity and optogenetic stimulation of these cells promotes sucrose consumption in sated mice (Garcia et al., 2021; Jennings et al., 2015), we investigated whether tesofensine would alter sucrose consumption induced by LH-GABAergic population activation in lean mice. To achieve this goal, we used sated transgenic Vgat-ChR2 mice (n=4) that constitutively expressed the opsin channelrhodopsin (ChR2) in GABAergic neurons to optostimulate in 5 minutes on and off blocks (Garcia et al., 2021). On different days, we injected saline, tesofensine 2 mg/kg + laser, or tesofensine 2 mg/kg alone without optostimulation (**Figure 4A**). The cumulative number of licks given in 30 minutes session can be seen in **Figure 4B**. Under saline injection, the Vgat-ChR2 mice increased sucrose intake selectively during optostimulation (see blue line). In contrast, under tesofensine treatment, the licks evoked by optostimulation of LH GABAergic neurons diminished (**Figure 4C**, see saline injection + laser vs. tesofensine+laser). These results indicate tesofensine attenuated open loop induction of sucrose feeding.

**Figure 4.**
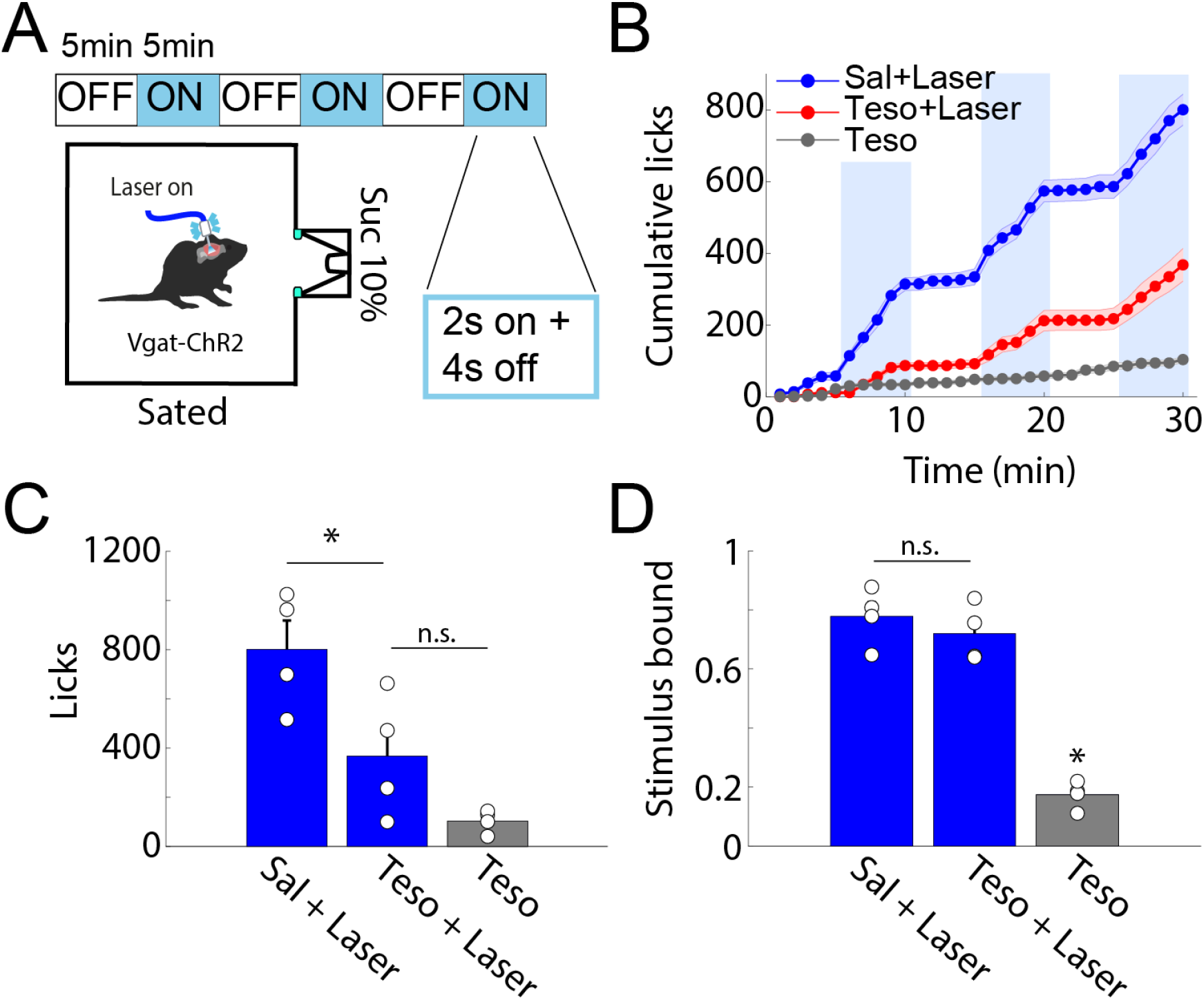
Tesofensine reduces the feeding behavior induced by the optogenetic activation of LH GABAergic neurons in lean Vgat-ChR2 mice. **A.** Schematic of the open-loop task, where VGAT-ChR2 mice (n = 4) with a fiber implanted in their LH, were stimulated at 50Hz for 2 seconds with 4 seconds off in blocks of 5 minutes (blue), while mice could lick a sucrose solution (10%) from a sipper. **B.** Cumulative licks during the open-loop stimulation with tesofensine (Teso) 2mg/kg or saline solution (Sal) administered by subcutaneous injection 30 minutes before the task. The tesofensine group (gray) was placed in the task, but the laser was not stimulated. Note that the teso+laser group exhibited an attenuated feeding response elicited by stimulation of LH GABAergic neurons. **C.** Total licks during a session, with blue bars representing the blocks that were laser stimulated in the task. **D.** Stimulus bound feeding that is the fraction of licks given during the stimulation window (spanning from the first laser pulse to 2.5 seconds after), with each circle representing a different mouse. Data are presented as mean ± SEM, and the results were statistically significant (\**p*-value < 0.05 based on paired t-test value).

In addition, it is well known that LH GABAergic stimulation typically leads to stimulus-bound feeding. Most feeding occurs within 2.5 seconds of optogenetic stimulation (Garcia et al., 2021; Valenstein & Cox, 1970) (**Figure 4D**; Sal + laser). In an open loop protocol (i.e., independently of behavior), we found that tesofensine treatment reduced the number of licks but did not affect stimulus-bound feeding (**Figure 4D**, Teso + Laser), showing that the drug per se did not impair oromotor reflexes elicited by optogenetic stimulation. These results demonstrate that the tesofensine-induced reduction in sucrose consumption, measured by the number of licks, is due to decreased feeding consummatory behavior rather than simply impairing oromotor reflexes elicited by optogenetic stimulation.

### The feeding response to optogenetic stimulation of LH GABAergic neurons was not significantly reduced by tesofensine 2 mg/kg in obese mice

After demonstrating the anorexigenic effects of tesofensine in lean Vgat-ChR2 mice, we aimed to replicate our findings in obese Vgat-IRES-cre mice. We expressed ChR2 in the LH through viral infection and exposed the mice to a high-fat diet or standard chow for 12 weeks (**Figure 5A**). We optogenetically stimulated LH GABAergic neurons in an open loop optogenetic stimulation paradigm and measured sucrose intake by drinking through a sipper filled with sucrose (**Figure 5B**). As a control, we injected an EYFP-expressing AAV vector. **Figure 5C** showed the cumulative licks in the Lean-EYFP control mice when saline, tesofensine 2mg/kg, or tesofensine 6 mg/kg was administered. Tesofensine 2 mg/kg significantly reduced the cumulative licks compared to saline, and the number of licks was further suppressed at the higher dose of 6 mg/kg, confirming the anorexigenic effects of tesofensine in mice (**Figure 5C**). In lean mice expressing ChR2 (Lean ChR2), sucrose intake increased within the optogenetic stimulation block, and both doses of tesofensine suppressed feeding evoked by optogenetic stimulation of LH GABAergic neurons (**Figure 5D**).

**Figure 5.**
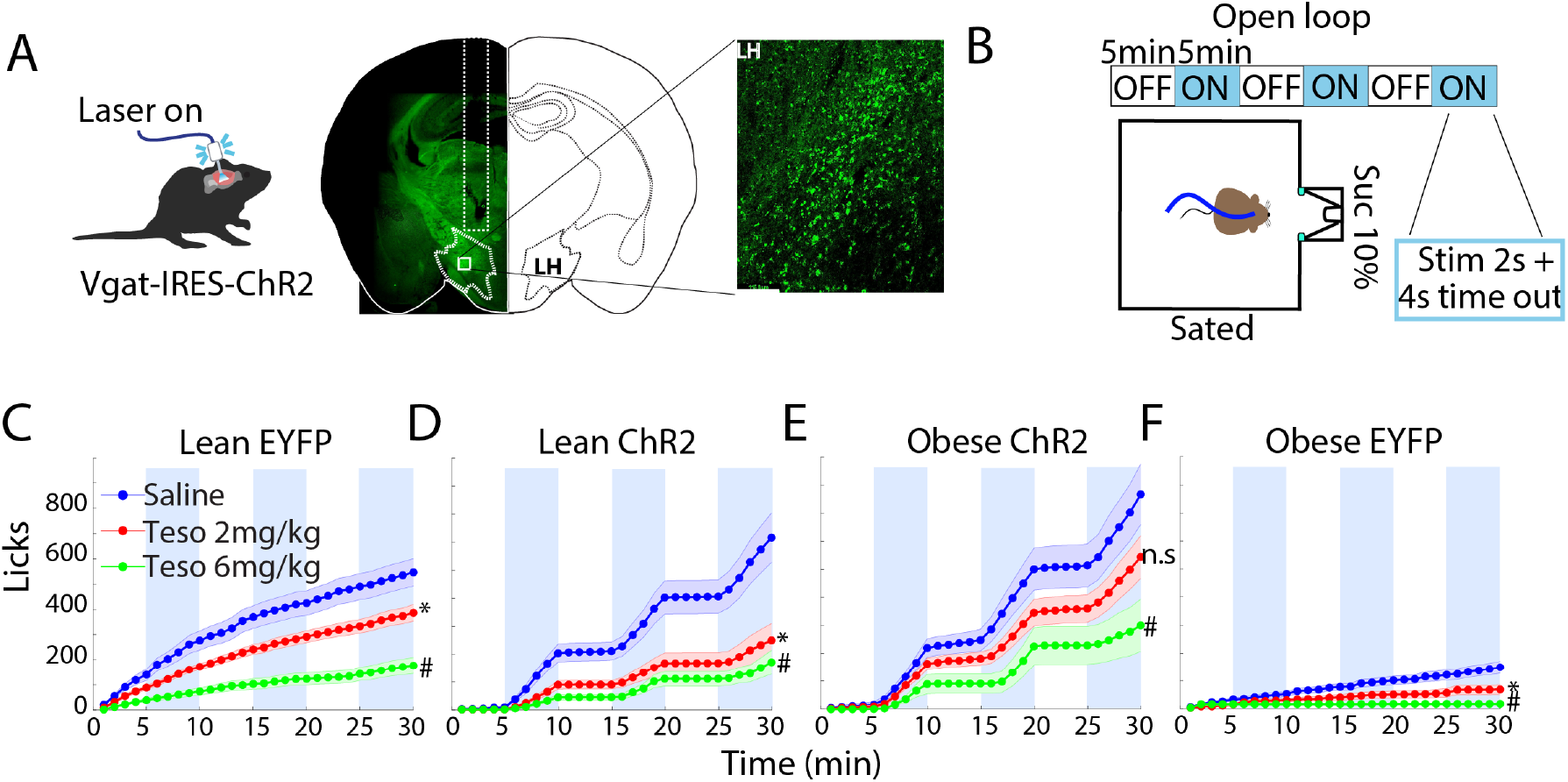
Tesofensine does not attenuate the optogenetic feeding elicited by LH GABAergic neurons in obese Vgat-IRES-cre mice. **A.** Representative image of Vgat-IRES-Cre mouse expressing ChR2 neurons (green) in the LH. **B.** Vgat-mice were fed with a HFD or standard Chow diet for 12 weeks, whereupon they were optostimulated. The mice had access to a sucrose solution (10%) in a sipper during the task. Mice had food and water *ad libitum* before the task. **C-F.** These four panels depict the cumulative licks (intake) during the open-loop task for lean and obese mice administered with saline, tesofensine (Teso) 2 mg/kg, or 6 mg/kg 30 minutes before the task. Data are presented as mean ± SEM. The results show that in obese mice tesofensine at 2 mg/kg did not reduce the cumulative number of licks during the open-loop stimulation protocol (**E**), as it did in lean mice (**D** see red lines). * *p* <0.05, Two-sample Kolmogorov-Smirnov test compared to control saline mice. # *p* <0.05 compared to tesofensine 2 mg/kg. n.s. p> 0.05 compared to saline obese ChR2.

In contrast, in obese ChR2-expressing mice (**Figure 5E**; Obese ChR2), tesofensine 2 mg/kg did not suppress optogenetically evoked feeding. Only the higher 6 mg/kg dose suppressed feeding (see # green trace). These results suggest that GABAergic neurons in obese mice are more efficacious to induce feeding than those in lean mice.

Paradoxically, we found that the control obese mice expressing EYFP (Obese-EYFP) consumed less sucrose overall compared to the Lean-EYFP mice (Two-sample Kolmogorov-Smirnov test, *p*<0.05), indicating that the HFD access may have indirectly led to sucrose devaluation (**Figure 5F**). We do not know the reason for this phenomenon, but similar phenomena have been previously reported (Beutler et al., 2020).

### The anorexigenic effects of tesofensine are amplified by the chemogenetic inhibition of LH GABAergic neurons

We then tested whether inhibiting LH GABA neurons could enhance the anorectic effects of tesofensine (2 mg/kg). To reach this end, we employed a FED3 system to enable mice to obtain chocolate pellets via nose-pocking in their homecages while chemogenetic inhibition was performed on the GABA neurons. We used the Vgat-IRES-cre mice to virally express hM4D(Gi) in the LH, an inhibitory DREADD (**Figures 6A, B**). Using this DREADD, the evoked action potentials are inhibited on clozapine-induced activation of hM4D(Gi)(Zhu and Roth, 2014). Clozapine-N-Oxide (CNO) infusion occurred at 18:30, and nose pokes and pellets delivered were continuously recorded for up to 24 hours. Our results revealed that the combination of Teso+CNO resulted in greater cumulative feeding suppression than any individual treatment during the first 19 hours post-treatment (**Figure 6C**, Kolmogorov-Smirnov test; Veh vs. CNO *p* = 0.006, CNO vs. Teso; *p* = 0.01, and Teso vs. Teso+CNO *p* =0.04). Thus, silencing LH GABAergic neurons enhanced the anorectic effects of tesofensine.

**Figure 6.**
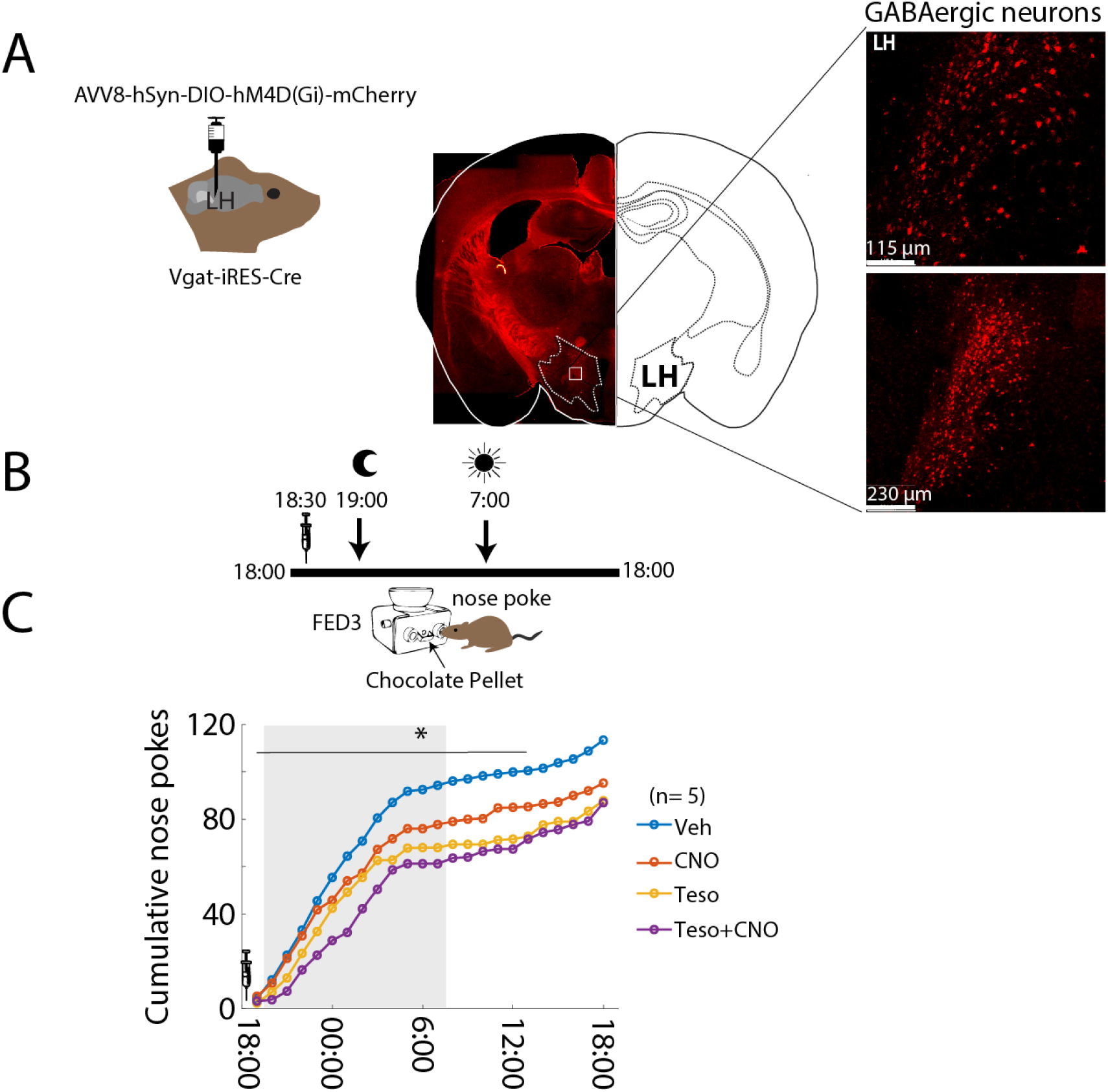
Chemogenetic silencing of LH GABAergic neurons potentiates tesofensine’s anorexigenic effects. **A.** Representative image of LH in mouse expressing hM4D(Gi), which inhibits GABAergic neurons chemogenetically. **B.** Schematic of a task where mice were placed in their home cages with an automated feeder (FED3) that delivered chocolate pellets for each nose poke. Recording started at 18:00 and lasted for 24 hours. At 18:30 h, the same five mice were injected with vehicle (Veh), Clozapine-N-Oxide (CNO) a ligand for hM4D(Gi), tesofensine (2 mg/kg), and CNO + tesofensine. The dark cycle lasted from 19:00 to 7:00 h. The moon symbol indicates when the ambient lights were turned off, and the sun when the lights were turned on. **C.** Average cumulative nose pokes during the 24-hour recordings for each group. The syringe indicates the time of drug administration at 18:30. The teso+CNO group performed fewer cumulative nose pokes and thus obtained fewer pellets than the other groups. * Significantly different Kolmogorov-Smirnov test (for the first 19 hours after drug administration see horizontal line); Veh vs. CNO *p* = 0.006, CNO vs. Teso; *p* = 0.01, and Teso vs. Teso+CNO *p* =0.04.

### Comparison with other appetite suppressants in lean rats

Having shown the neuronal correlates of tesofensine in the LH in rats and mice, we compared tesofensine appetite suppressant effects with other appetite suppressants, particularly phentermine and 5-HTP.

### Tesofensine induces body weight loss in rats without producing head weaving stereotypy at therapeutic doses

It has been proposed that tesofensine has an important dopaminergic component (Appel et al., 2014; Bara-Jimenez et al., 2004; Hansen et al., 2013). Hence, the motor effects of tesofensine were compared against phentermine, a hallmark dopamine-acting appetite suppressant. Involuntary movements with no apparent function are named stereotypies (Asser and Taba, 2015; Kelley, 1998). Our research group recently reported that head weaving stereotypy is a common side effect of most appetite suppressants, particularly those acting to enhance DA efflux, such as phentermine (Kalyanasundar et al., 2015; Perez et al., 2019). Head weavings were indicated by unnecessary head oscillations. Therefore, we characterized the tesofensine-induced stereotypy effects compared with phentermine, an amphetamine congener that served as a positive control. To quantify stereotypic behavior, we used DeepLabCut, a markerless pose estimation tool based on transfer learning with deep neural networks (Mathis et al., 2018). We trained the network to detect a rat’s nose, forelimbs, and tail base from a bottom-view videotaped session (see **Video 1**). The video recording lasted for four hours after the drug administration. We observed that the control rats treated with saline exhibited a physiological level of forward locomotion (**Figure 7A**). Likewise, they spent about 65% of the session in a quiet-awake state (refer to **Video 1**), most often in a "sleeping" position (**Video 2**), which we pooled together for analysis (**Figure 7B**). Our algorithm incorrectly identified "head weaving stereotypy" in control rats, as these animals did not exhibit this behavior. This is because our algorithm identified a part of the grooming sequence and misclassified it as stereotypy (refer to **Video 3** and (Berridge et al., 1987)), likely because grooming and head weaving share certain similarities (**Figure 7C**). Nonetheless, this “grooming” behavior occurred randomly with low probability (**Figure 7C**; Vehicle, i.p.) and with variable onset times (**Figure 7D**).

**Figure 7.**
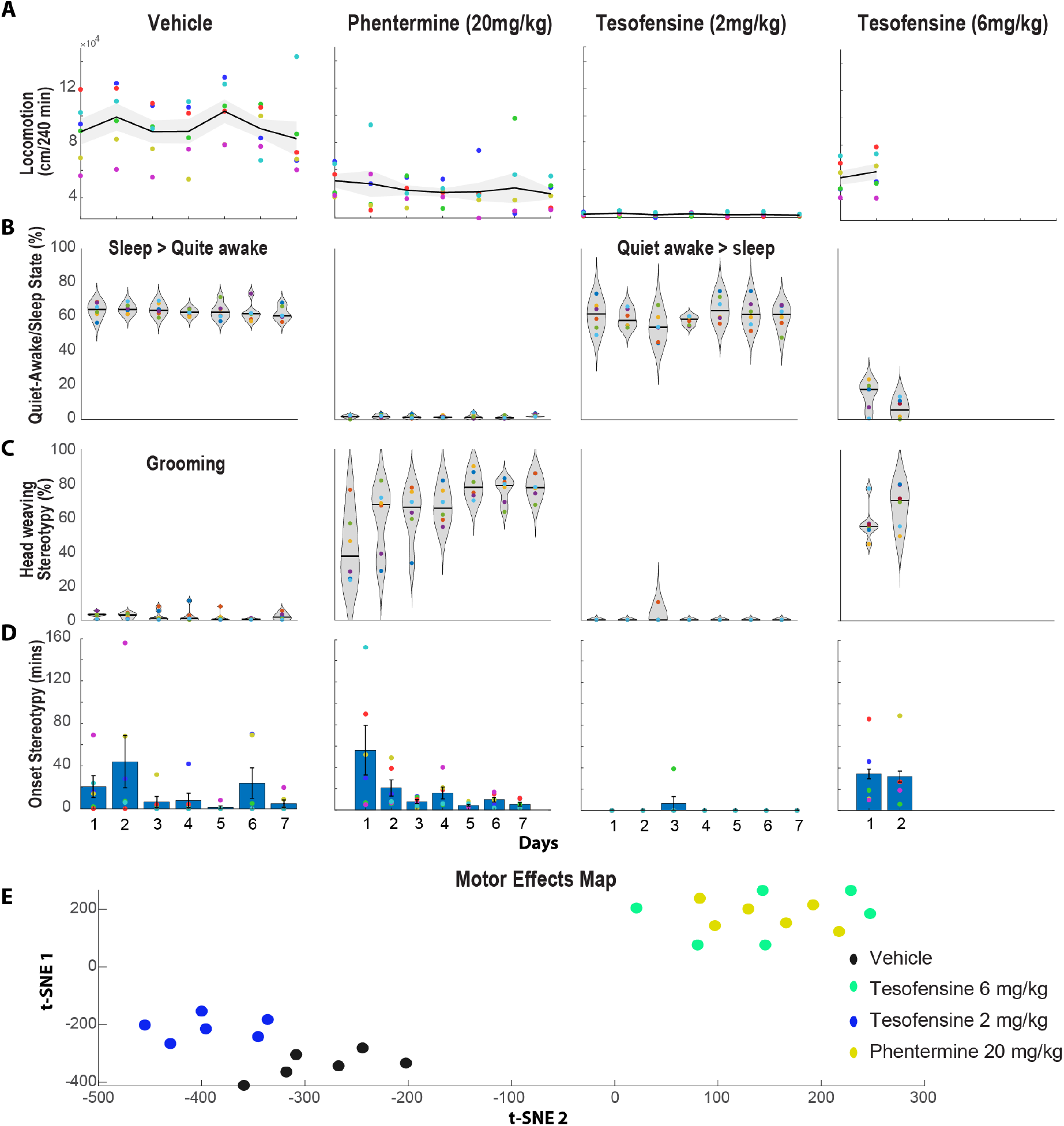
Effects of tesofensine and phentermine on locomotion, quiet-awake/sleep, and head weaving stereotypy in rats. **A.** Total distance traveled (cm/240 min) during forward locomotion over seven days of treatment with saline (1 ml/kg, n= 6), phentermine (20 mg/kg, n= 6), tesofensine (2 mg/kg, n= 6), or tesofensine (6 mg/kg, n= 6). Each point depicts one rat, with the black line indicating the mean and the gray shaded area indicating the standard error of the mean of the distance traveled by all rats in the group. All treatments reduced locomotion **B.** The proportion of time that rats spend in a quiet-awake/sleep state is defined as when the rat is not moving but either awake or sleeping. Only the Vehicle and Tesofensine 2 mg/kg groups spent more than 60% of their time in these behavioral states. **C.** The distribution of the percentage of time that rats exhibited a head weaving stereotypic behavior over seven days following treatment injection. Each point in the plot represents one rat, and the width of the violin plot indicates the probability density distribution. The black line shows the median value for each day. For phentermine, the results showed that the percentage of time the rat spent exhibiting head weaving stereotypy gradually increased across the seven days. Additionally, head weaving stereotypy was aggravated across days in all rats treated with phentermine. In Vehicle control rats, grooming behavior was mistakenly classified as head weaving stereotypy. This is because control rats do not express this behavior. Surprisingly, rats treated with tesofensine 2 mg/kg did not exhibit much stereotypy, and thus apparently neither grooming. **D.** The onset of the first event of stereotypy was measured across days. Note that for phentermine the onset of stereotypy decreases across days. **E.** For the first 2 days, the variables locomotion, quiet awake/sleep, onset, and stereotypy were analyzed using a clustering algorithm (t-SNE). This analysis uncovered two main clusters: the first corresponds to rats treated with vehicle and tesofensine 2 mg/kg. Note that rats treated with tesofensine 2 mg/kg were in a slightly different position than control rats. The second cluster mixed rats treated with phentermine and tesofensine 6 mg/kg. Hence, t-SNE seems to separate rats according to their overall motor profile effects induced by each drug. Overall, at therapeutic doses, tesofensine induced body weight loss without producing head weaving stereotypy.

Phentermine via i.p. resulted in a slightly increased locomotion and decreased time spent in a quiet-awake/sleep state (**Figures 7A-B**; Phentermine). Interestingly, DeepLabCut analysis unveiled for the first time that phentermine-treated rats exhibited less forward locomotion than control rats (despite it being a stimulant drug; **Figure 7A**). Notably, phentermine induced strong head weaving stereotypy, which increased gradually over seven days and occupied 80% of the time of the 4-hour session (**Figure 7C**). Head weaving stereotypic behavior involved rats standing still on four legs and moving their head erratically (**video 4**), accompanied by frequent uncontrolled tongue movements (although we did not formally quantify tongue movements, we report them as a subjective human visual observation). The onset of stereotypy decreased from 56.1 ± 23.2 minutes on the first day to 5.5 ± 1.8 minutes on the seven days of treatment (**Figure 7D**).

In contrast, at a low dose of tesofensine (2 mg/kg) induced little or no forward locomotion (**Figure 7A**). Rats spent more time in a quiet-awake state than in a sleep position (**Figure 7B**), and head weaving stereotypy was detected in only one rat and for a short period (**Figure 7C**; day 3, **Video 5**). As noted, our algorithm in control rats erroneously misclassified grooming behavior as stereotypy in control rats. However, no head weaving stereotypy was detected under tesofensine 2 mg/kg, suggesting, at least indirectly, a decrease in the probability of grooming behavior. Nevertheless, in rare instances, we observed that rats in a quiet-awake state would also execute jaw and tongue movements, albeit at a lower intensity (see **Video 6**). Further studies are needed to evaluate these effects more carefully.

Finally, a high dose of tesofensine (6 mg/kg) was administered for two days only to avoid lethality, which led to increased locomotion and reduced time spent in a quiet awake/sleeping state (**Figures 7A-B**). At this high dose, rats exhibited clear and robust stereotypy behavior with rapid onset (**Figures 7C-D**), primarily comprising uncontrolled tongue movements and less intense head waving (**Video 7**). From a visual inspection, we note that the stereotypy induced by tesofensine differs slightly from that induced by phentermine. However, both drugs share the common feature of inducing uncontrolled tongue movements, which earlier studies had failed to report. In summary, tesofensine at a low dose induced almost no head weaving stereotypy, but a robust stereotypy was observed at a high dose.

The motor map effects were represented using t-SNE (**Figure 7E**). This algorithm clusters rats’ behavior based on their overall profile of changes in motor variables, including locomotion, quiet awake/sleep time, onset, and stereotypy. Each dot on the map is a rat, and rats with similar profiles are grouped together. We observed that rats treated with tesofensine 2 mg/kg exhibited different behavior compared to the control group. In contrast, rats treated with tesofensine 6 mg/kg and phentermine, which both exhibited more stereotypy, were grouped in a small area but far away from the rats in the control and tesofensine 2 mg/kg groups (**Figure 7E**). Further studies are needed to investigate the effects of tesofensine on reducing the likelihood of grooming behavior and other tongue kinematics parameters.

### Pharmacological interaction with a serotonin appetite suppressant

#### Tesofensine prolongs the body weight loss effect of 5-HTP/CB, a serotonin precursor

In lean animals, we evaluated the weight loss effects and interaction of tesofensine with 5-HTP, another appetite suppressant, by administering each drug alone or in combination. The change in body weight (g) after administration of control (vehicle), tesofensine (1 and 2 mg/kg), 5-HTP (31 mg/kg), and CB (75 mg/kg) is shown in **Figure 8A**. We found a significant difference in body weight among groups [RM ANOVA; main effect doses: F(5,28)= 26.4, *p* < 0.0001; days: F(14,392)= 22.1, *p* < 0.0001; doses per days interaction: F(70,392)= 6.7, *p* < 0.0001]. The control group steadily gained body weight, while tesofensine (1 mg/kg) prevented body weight gain without achieving statistical differences compared to control rats (*p* > 0.3, n.s.). At 2 mg/kg, tesofensine induced modest but significant weight loss from the eighth day of treatment onwards (*p* < 0.05). 5-HTP/CB caused rapid, substantial weight loss for the first six days of treatment but showed gradual weight regain, indicating tolerance. Combining both drugs led to substantial weight loss from 1 to 8 days, which was maintained until the end of treatment. The combination led to greater weight loss than each drug alone (*p* < 0.05). Our results demonstrated that tesofensine reverted the pharmacological weight loss tolerance observed in 5-HTP/CB alone.

**Figure 8.**
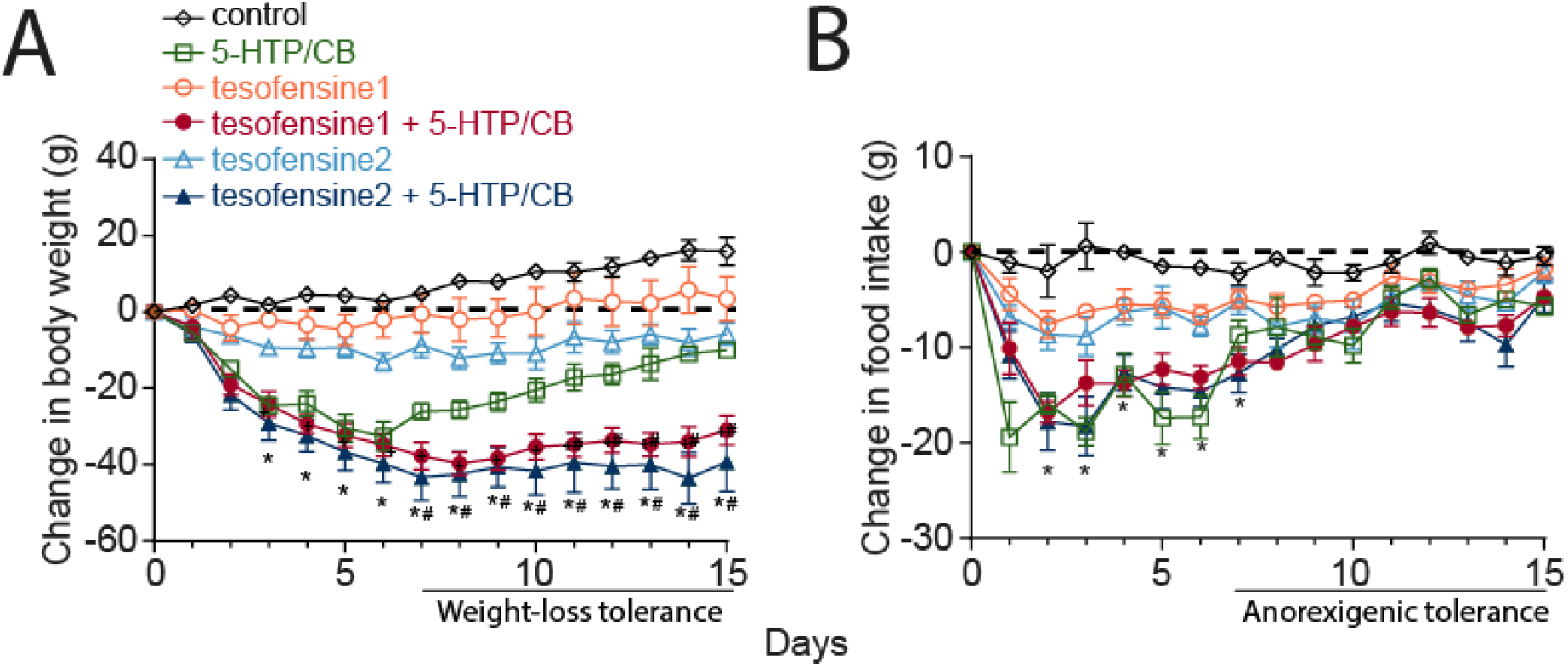
Tesofensine prolongs body weight loss produced by serotonin precursor 5-HTP/CB on the chow control diet. **A.** Change in body weight (g) relative to pretreatment day 0. Tesofensine1 and tesofensine 2 refer to doses of 1 and 2 mg/kg administered subcutaneously, respectively. 5-HTP/CB refers to doses of 31 and 75 mg/kg administered intraperitoneally. Note that the triple combination of tesofensine, 5-HTP, and CB led to significantly greater weight loss than any other group and did not show weight loss tolerance as seen in 5-HTP/CB group (see horizontal line at days 7-15). **B.** Changes in food intake for the groups shown in panel “**A**.” Note that 5-HTP/CB, tesofensine1, and 2 + 5-HTP/CB groups exhibited anorexigenic tolerance, as evidenced by their increased food intake from day 7 to 15. Data are presented as mean ± SEM. \**p* < 0.05 compared to tesofensine2; +*p* < 0.05 compared to tesofensine1; #*p* < 0.05 compared to the 5-HTP/CB group.

Regarding food intake, **Figure 8B** shows the change in food intake that accompanied weight loss. The control group showed no change in food intake (−1 ± 0.2 g). Tesofensine (1 and 2 mg/kg) reduced food intake (−4.7 ± 0.4 g and -6.1 ± 0.5 g, respectively). A more potent anorexigenic effect was induced by 5-HTP/CB alone. 5-HTP/CB alone suppressed food intake for 1-6 days (−16.9 ± 0.9 g) but gradually increased the intake for 7-15 days (−6.6 ± 0.7 g). A similar tolerance to the anorexigenic effects of the combination was also seen. Both combinations, tesofensine1 or 2, + 5-HTP/CB suppressed food intake from 7 to 15 days (−8.1 ± 0.7 g and -7.8 ± 0.8 g, respectively). However, they demonstrated no synergistic effect compared to 5-HTP/CB alone (p > 0.1, n.s.). Hence, the tolerance for anorectic effects was still present in the combination. However, tesofensine entirely reverted the 5-HTP tolerance for weight loss, suggesting a greater effect of tesofensine on energy expenditure than food intake suppression.

#### The tesofensine plus 5-HTP/CB combination did not achieve a significant synergy in a sucrose intake suppression on an isobolographic assay

To analyze the effect of tesofensine plus 5-HTP/CB, we evaluate the consumption of oral sucrose. **Figure 9A** shows the percentage of the anorexigenic effect induced by different doses of tesofensine. The administration of tesofensine increases the anorexigenic effect in a dose-dependent manner, demonstrating that 0.71, 1.9, and 5 mg/kg were significant comparisons to the control group (*p* < 0.05, see *). Tesofensine at 5 mg/kg induced the maximum suppressed sucrose intake (51.2 ± 5.1 %), while 0.1 mg/kg did not suppress it (−6.3 ± 9.3 %). The 5-HTP/CB dose-dependently decreased sucrose consumption at 1.8, 7, 26.6, and 100 mg/kg (*p* < 0.05 to the control group; **Figure 9B**). 5-HTP/CB at a 100 mg/kg dose achieved the maximum suppressed consumption (98.3 ± 0.4%). Regarding tesofensine, the ED30 value was 2.11 ± 0.8 mg/kg, while 5-HTP/CB had an ED30= 6.36 ± 2.2 mg/kg (**Table 1**). Then, we used the ED30 values of tesofensine and 5-HTP to evaluate the combinations. Tesofensine + 5-HTP/CB at 1:1 (0.5, 1, 2.1, and 4.2 mg/kg), and at 3:1 (0.3, 0.7, 1.5, and 3.1 mg/kg) proportion decreased sucrose intake at a dose-dependently manner (**Figures 9C, E**). Isobolographic analysis showed that the experimental ED30 (red diamond) was lower than the theoretical ED30 (**Figures 9D, F**), suggesting a tendency for a synergistic effect. However, using Tallarida’s t-student test (Tallarida, 2000) revealed no significant differences between the experimental and theoretical effective doses in the 1:1 and 3:1 proportions (**Table 1**). Therefore, our isobolographic analysis suggests no significant pharmacological synergism between tesofensine and 5-HTP, at least in our acute sucrose suppression assay.

**Figure 9.**
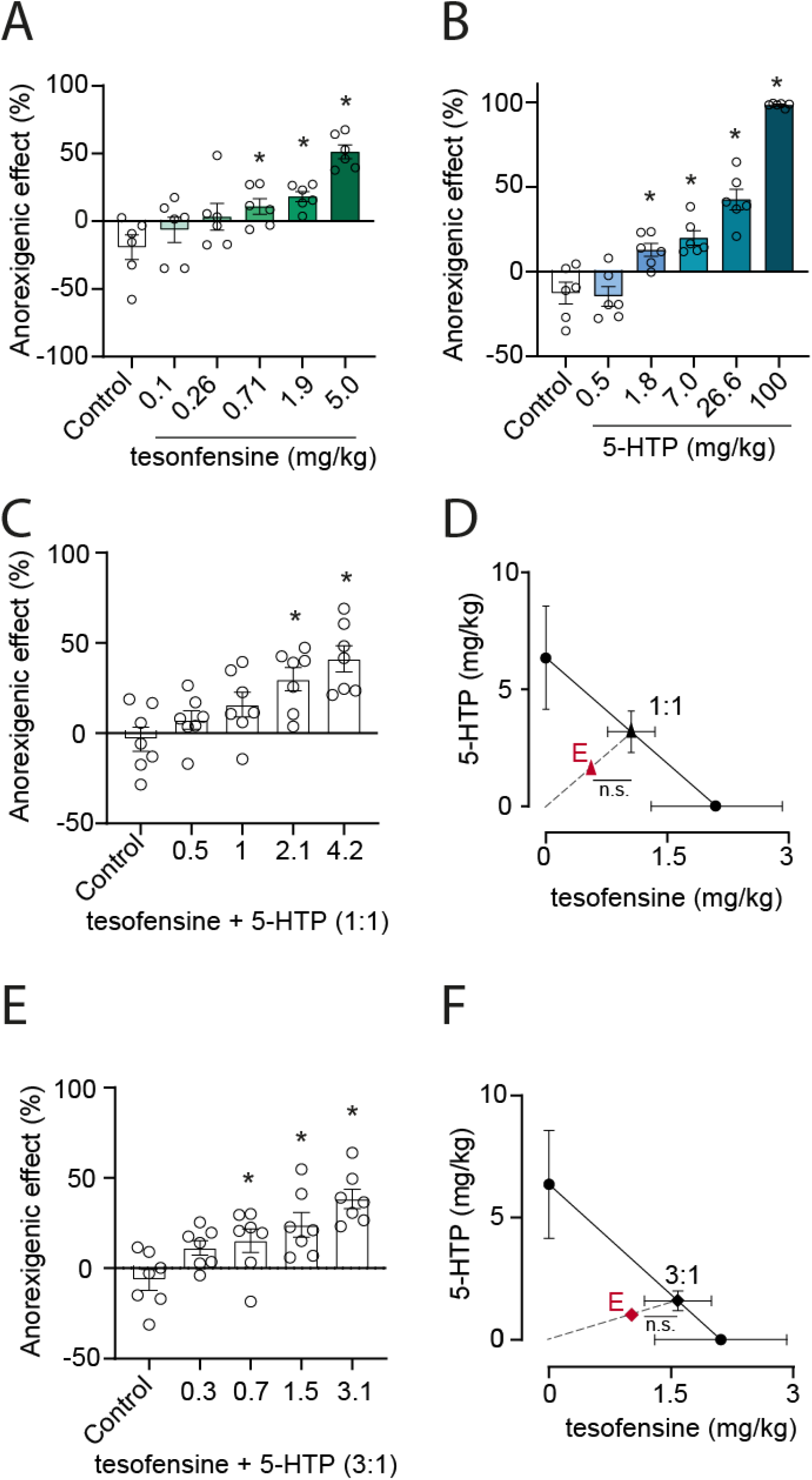
In rats, the combination of tesofensine and 5-HTP is not more effective than either drug alone in reducing sucrose intake. Panel **A** shows the dose-response curve for tesofensine, and panel **B** shows the dose-response curve for 5-HTP. The curves show that both tesofensine and 5-HTP reduce sucrose intake in a dose-dependent manner (i.e., the % anorexigenic effect). See **Figure 9 Supplementary 1** for sucrose intake data before, during, and after drug administration. **C.** The anorexigenic effect of the tesofensine + 5-HTP combination at a 1:1 ratio. **D.** Isobologram graph depicting the anorexigenic interaction at a 1:1 ratio. The x and y axes points indicate the experimental ED30 values of tesofensine and 5-HTP, respectively. At the same time, the red diamond (letter **E**) represents the ED30 obtained experimentally for the combination at a 1:1 ratio. The diagonal line connecting the ED30 of tesofensine and 5-HTP represents the theoretical line of additivity, and the point on this line represents the theoretical ED30 of the combination at a 1:1 ratio. The results show that the combination of tesofensine and 5-HTP is not significantly more effective than either drug alone. **E.** Anorexigenic effect of the tesofensine + 5-HTP combination at a 3:1 ratio. **F.** Isobologram graph indicating the anorexigenic interaction at a 3:1 ratio. The same conventions as panel **D**. Data are presented as mean ± SEM. \**p* < 0.05 compared to the control group. n.s. indicates no statistical difference between the experimental ED30 dose and the theoretical ED30.

**Table 1.** Effective dose 30 obtained by linear regression of the dose-response curve of tesofensine and 5-HTP.

| Drug | ED30 ± SEM | Pearson correlation coefficient (r) |
| --- | --- | --- |
| Tesofensine | 2.11 ± 0.81 mg/kg | 0.874 |
| 5-HTP | 6.36 ± 2.21 mg/kg | 0.906 |

| Statistical analysis of tesofensine + 5-HTP interaction |  |  |  |
| --- | --- | --- | --- |
| Tesofensine + 5-HTP/CB | Theoretical ED30 ± SEM | Experimental ED30 ± SEM | Interaction index |
| 1:1 | 4.24 ± 1.1 mg/kg | 2.21 ± 0.1 mg/kg (n.s.) | 0.52 |
| 3:1 | 3.17 ± 0.8 mg/kg | 2.03 ± 0.3 mg/kg (n.s.) | 0.64 |

ED30= effective dose 30; SEM= Standard Error of the Mean (SEM), the results were not statistically significant (n.s.) from the theoretical ED30.

One probable reason for the appetite-suppressing effect of tesofensine (or 5-HTP) is that it may induce taste aversion. However, our study results do not support this hypothesis. As shown in **Figure 9 Supplementary 1**, the sucrose consumption levels almost returned to baseline after the injection of 5-HTP (**Figure 9 Supplementary 1A**) or tesofensine (**Figure 9 Supplementary 1B**) on the next day (day 8). This suggests that taste aversion is unlikely to be the primary mechanism behind the anorexigenic effect of these appetite suppressants.

#### Tesofensine does not affect sucrose detection or oromotor palatability responses

To investigate whether tesofensine impairs sucrose detection or palatability responses, we modified a sucrose detection psychophysical task using a new equipment called the homegustometer (Fonseca et al., 2020, 2018). Rats performed the task day and night over multiple consecutive days, which allowed us to characterize the temporal pharmacological effects of tesofensine across (**Figure 10A**) and within days (**Figure 11**). The rats (n=4) were trained to discriminate between different concentrations of sugar and water using a homegustometer, and their performance was recorded continuously for up to 23 hours daily. At the start of each trial, rats visited the central port and delivered 2-5 dry licks, triggering a taste stimulus, including a single drop of water or one of five sucrose solutions with varying concentrations. The rats discriminated water from sucrose by moving leftward for water and rightward for sucrose solutions (counterbalanced between subjects), with successful discrimination resulting in a reward (three drops of water in lateral outcome sippers). After the rats learned the sucrose detection task, we injected them with 2 mg/kg of tesofensine at ∼ 18:00 h for five days. We observed that tesofensine did not impair daily task performance (**Figure 10B**; %Correct). The number of trials and thus total consumption, we observed a temporal increase in the first and second days of baseline that gradually reached an asymptote (**Figure 10B**; Trials and Consumption). Likewise, we observed that psychometric curves during the baseline, tesofensine, and post-tesofensine days were practically identical (**Figure 10C**). Although we also note that under and after tesofensine treatment, the subjects tend to be more sensitive to sucrose, detecting it correctly even at the lowest sucrose concentration (0.5%) (**Figure 10C**). Thus, it is unlikely that the tesofensine anorexigenic properties are due to diminished sweetness detection or changes in their hedonic value or palatability.

**Figure 10.**
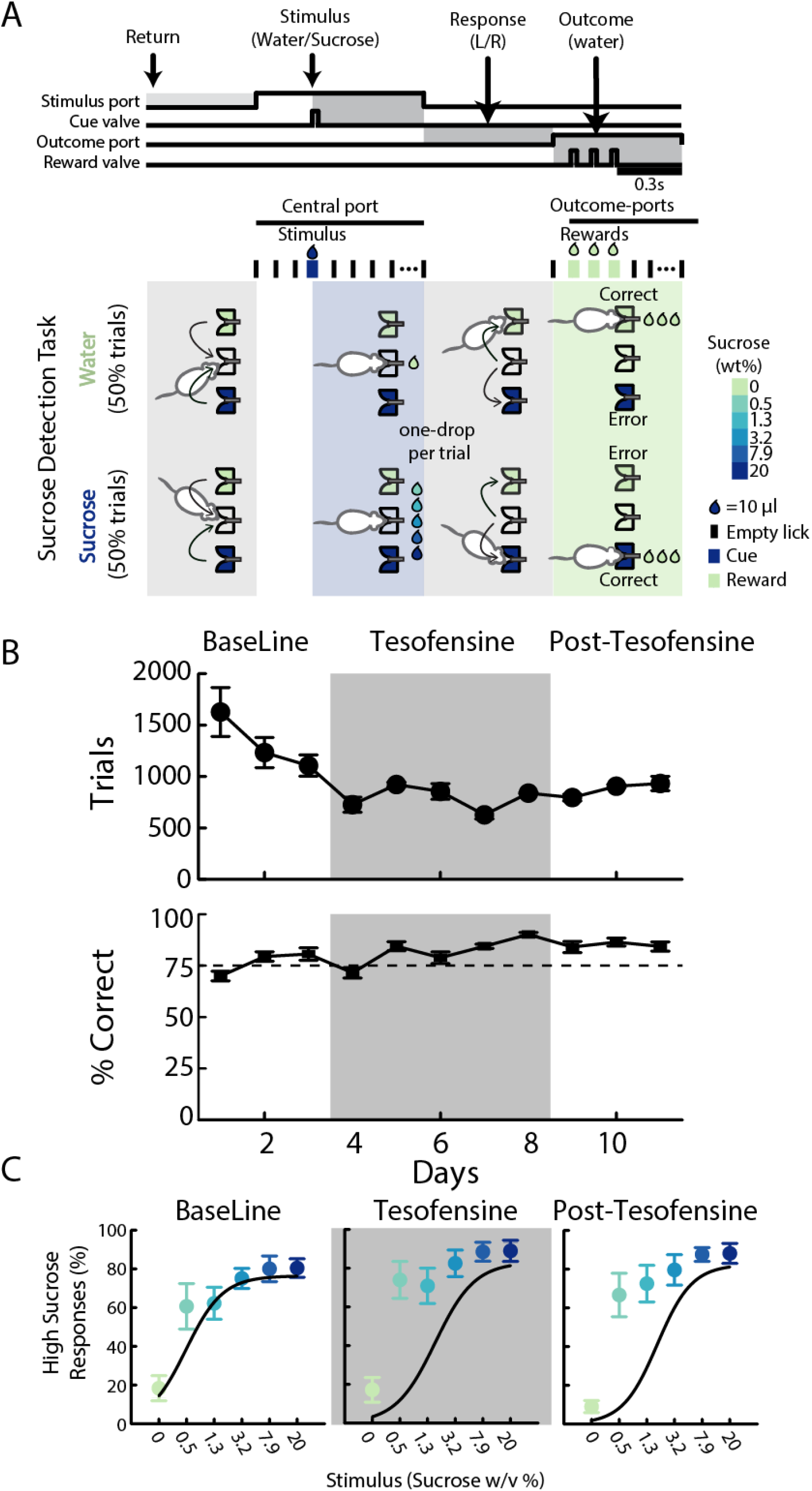
Tesofensine did not affect performance of rats in a sucrose detection task. **A.** Rats were trained to lick a central spout that dispensed the stimulus a drop of water or solutions of sucrose. To obtain a reward (3 drops of water), rats had to choose between two lateral spouts. **B.** Upper panel shows the number of trials, and the lower panel the correct performance across the baseline, tesofensine treatment, and post-tesofensine days. There were no significant differences in the percent correct, the trials per session, or the total volume consumed between these periods, except for an overall decrease in the number of trials during the baseline period as the rat re-learned the task. The gray rectangle depicts days of tesofensine administration **C.** Plots of high sucrose responses as a function of sucrose concentration. The psychometric curves for the sucrose detection task also did not differ significantly between the baseline, tesofensine, and post-tesofensine periods. These findings suggest that tesofensine does not affect performance in the sucrose detection task in rats. The x-axis is scaled logarithmically, and the data are represented as mean ± SEM.

**Figure 11.**
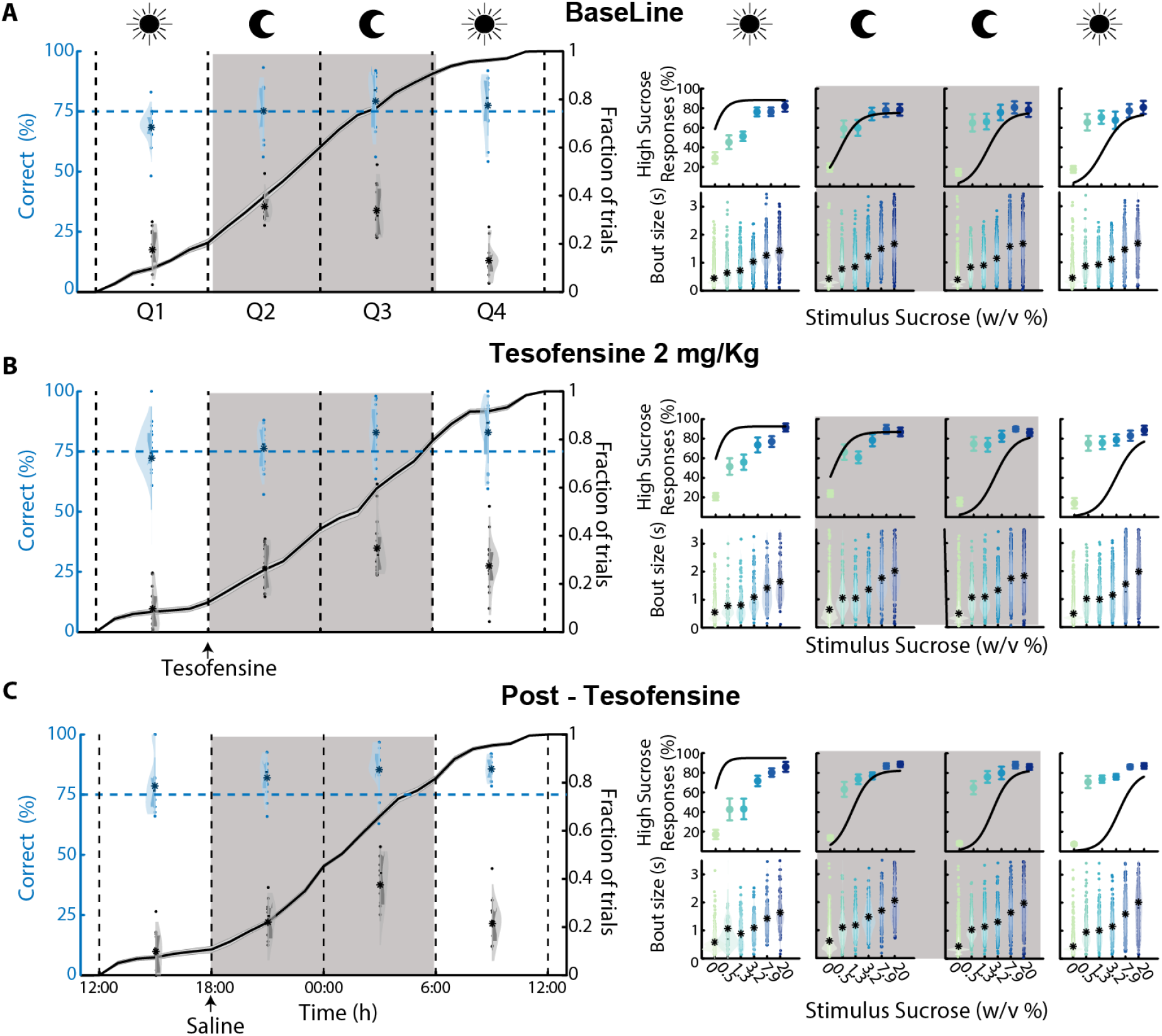
Performance of rats in a sucrose detection task across 23 hours under baseline, tesofensine, and post-tesofensine treatments. **A.** Baseline performance. The data are presented in 6-hour intervals (divided into four quartiles, Q1–Q4-the moon symbol indicates quartiles during night and sun symbol periods of light) and include the percentage of correct responses and the fraction of trials performed in each quartile across the days. The chance is 50% for the left blue axis. **B**. Tesofensine treatment (2 mg/kg). The data are presented in the same way as in panel “**A**.” **C.** Post-tesofensine treatment. For all panels, the psychometric curves at the right show the percentage of correct choices for detecting sucrose solutions as a function of sucrose concentration. The x-axis of the psychometric curve is scaled logarithmically. Below, the bout size is the time elapsed between the last lick in the central port after stimulus delivery. * Depicts the mean bout size, and each color dot is one single trial. The larger the bout size, the more palatable the drop of solution is to the rat. Our results indicate that sucrose detection and palatability responses were unaffected by tesofensine. The data are represented as mean ± SEM.

### Sucrose detection within a single day

This study investigated the effects of tesofensine administration on the perception of sugar taste in rats over a day. The data were analyzed by dividing the day into quartiles (6-hour periods) to investigate whether tesofensine affected the rats’ perception of the sucrose taste. First, we discuss the performance of the task under control baseline conditions (**Figure 11A**). As expected, given that rats are nocturnal animals, our results showed that they were more likely to complete more trials during the second and third quartiles of the day (Q2 18:00 to 00:00 - and Q3 00:00 to 6:00), corresponding to the dark cycle of the day. 30.5% of all trials were completed during the daytime (Q1, Q4), while 69.3% were performed at night (Q2, Q3, **Figure 11A**; see black percentages). However, the accuracy of the sucrose detection task was not significantly different across the day.

However, rats exhibited a non-significant tendency to decrease performance in Q1 from 12:00 to 18:00 relative to the other quartiles. This poor performance could explain the difficulty of fitting a psychometric curve during Q1 (see **Figure 11A**, right upper panels). The oromotor palatability responses evoked by the delivery of one drop of water or sucrose revealed a gradual increase in the bout size as a function of sucrose, reflecting the hedonically positive reaction that sucrose evokes (Perez et al., 2013). In addition, the palatability responses also did not vary across the day (see **Figure 11A**, right lower panels).

The major change observed during tesofensine treatment was a shift in the distribution of trials completed on each quartile. Specifically, rats performed significantly fewer trials in Q1 and Q2 but compensated for this by performing significantly more trials in Q3 and Q4. Thus, tesofensine treatment shifted the distribution of trials toward the right. However, the accuracy of the sucrose detection task (i.e., the percent correct trials) was not significantly altered by tesofensine (**Figure 11B**).

Finally, in the post-tesofensine period, rats received subcutaneous injections of saline. Given that the half-life of tesofensine is about 8 days, we continued evaluating the rats’ performance for three more days (**Figure 11C**). We observed no major change in task performance, or the palatability responses sucrose elicited during this period. Our data suggest that tesofensine in rats did not impair sweetness detection or affect its palatability.

## DISCUSSION

This study found that tesofensine induced greater weight loss in obese rats than in lean Wistar rats. We hypothesized that this was due to tesofensine’s ability to modulate neuronal activity in the LH. Our electrophysiological results showed that tesofensine produced a stronger and larger modulation of LH ensemble activity in obese rats than in lean rats. This suggests that tesofensine may act, in part, by modulating neuronal activity in the LH to reduce food intake and promote weight loss. More importantly, we also found that tesofensine inhibited GABAergic neurons in the LH of Vgat-ChR2 and Vgat-IRES-cre transgenic mice. These neurons promote feeding behavior optogenetically(Garcia et al., 2021; Jennings et al., 2015), so the inhibition of these neurons by tesofensine may contribute to its appetite-suppressing effects. In addition to its effects on the LH, in rats, tesofensine did not produce head weaving stereotypy at therapeutic doses, suggesting that it may be a safer and more tolerable option to treat obesity than other appetite suppressants such as phentermine. It also did not significantly potentiate the acute suppression of sucrose intake induced by 5-HTP, but it prolonged the weight loss induced by 5-HTP, a serotonin precursor and appetite suppressant. This suggests that tesofensine may be a useful adjunct to serotoninergic agents to treat obesity. Finally, we found that the appetite suppressant effect of tesofensine is not due to the induction of taste aversion. Rats resumed drinking sucrose right after the next treatment day in the isobolographic assay. Further studies using a 23-hour psychophysical sucrose detection task also showed that tesofensine might not affect the perception of sweetness or its palatability responses, even though it is a weight-loss drug. Taken together, our study provides new insights into the effects of tesofensine on weight loss and the underlying neuronal mechanisms. These findings suggest that tesofensine may be a promising new therapeutic agent for the treatment of obesity.

### Tesofensine targets the LH, silencing a subset of GABAergic neurons

Tesofensine is more efficacious in inducing weight loss in obese rats than lean Wistar rats. Our results replicate and confirm the findings observed by Hansen et al., 2013 in Sprague Dawley rats and (van de Giessen et al., 2012) in obese Wistar rats, suggesting that this is a robust attribute of tesofensine. They suggested that the grater efficacy was due to the ability of tesofensine to restore lower DA levels in the nucleus accumbens observed in obese rats (Hansen et al., 2013). Here, we further extend the neuronal correlates to the LH and uncovered for the first time that tesofensine produced a stronger and larger modulation of LH ensemble activity in obese rats than in lean rats. The effects of tesofensine on LH activity were complex. However, tesofensine seems to enhance the recruitment of LH neurons exhibiting activation after drug administration (i.e., see E4 neurons in **Figure 2**). The identity of this cell type is out of the scope of this study, but it is tempting to speculate that most likely includes a large subset of non-GABAergic neurons, perhaps enriched of glutamatergic neurons. We acknowledge that our data cannot rule out the intriguing possibility that a different subset of GABAergic neurons (from those inhibited) could be activated by tesofesnine. This is because activation of GABAergic neurons can trigger oromotor stereotypy (Nieh et al., 2015), similar to that observed with phentermine and tesofensine at high concentrations (see below **Figure 7**). Further studies using Cal-light or TRAP-like techniques should be conducted to confirm the identity of the activated neuronal ensembles recruited by tesofensine (Guenthner et al., 2013; Lee et al., 2017). These techniques could capture functional ensembles, enabling more precise identification of the cells that respond to tesofensine and are responsible for its therapeutic anorexigenic effects and stereotypies side effects.

The second larger group of cells that were more strongly modulated by tesofensine in obese than in lean rats was the ensemble of neurons exhibiting a robust inhibition (see E1 in **Figure 2**). Our data in Vgat-IRES-cre mice demonstrate that these neurons correspond to a subset of LH GABAergic neurons (**Figure 3**). We uncovered that tesofensine could silence a subset of optogenetically identified LH GABAergic neurons using optrode recordings. It also impaired their ability to be activated by an open loop optogenetic stimulation (**Figure 3**). Using lean Vgat-ChR2 mice, we found that tesofensine reduces the feeding behavior induced by the optogenetic activation of LH GABAergic neurons (**Figure 4**). Moreover, in Vgat-IRES-cre obese mice, only a higher tesofensine dose could suppress optogenetically induced feeding, suggesting that, during obesity, LH GABAergic neurons seem to be hypersensitized. Thus, these neurons are more resilient to evoked feeding during obesity (**Figure 5**). Conversely, the chemogenetic inhibition of LH GABAergic neurons potentiates the anorexigenic effects of tesofensine (**Figure 6**). Our data is the first to demonstrate that tesofensine directly targets LH feeding circuits, particularly silencing a subset of GABAergic neurons, and activating a still unidentified cell type (perhaps a subset of glutamatergic neurons). It paves the way to uncover better ways to enhance the therapeutic effects of tesofensine and perhaps for other appetite suppressants.

Recently, tesofensine has demonstrated promising results for treating rare human feeding disorders, such as hypothalamic obesity (Huynh et al., 2022). Hypothalamic obesity symptoms include exacerbated hunger, rapid increase in body weight, and low metabolism. Approximately 50% of craniopharyngioma survivors develop hypothalamic obesity (Müller et al., 2004). This type of tumor most often affects the physiological function of the hypothalamus, a part of the brain that regulates appetite and metabolism, thus leading to rapid, intractable weight gain, a condition known as hypothalamic obesity (Müller et al., 2004). In particular, the lack of satiety feedback from the hypothalamus has been proposed as a mechanism for hypothalamic obesity (Bray et al., 1981; Brobeck, 1960, 1946). Hypothalamic obesity is a challenging condition to treat, as there are currently no approved or effective pharmacological treatments. However, tesofensine is a novel compound with potential in human studies and may be a promising alternative for these patients (Huynh et al., 2022). Given the ability of tesofensine to modulate the activity of the LH, our preclinical findings agree with the proposal that tesofensine could be a useful treatment for patients with hypothalamic obesity, a rare feeding disorder, as recently demonstrated (Huynh et al., 2022).

### Comparison of tesofensine with other appetite suppressants

#### Tesofensine vs. a dopaminergic agent: phentermine

Tesofensine was initially intended to treat Parkinson’s and Alzheimer’s disease because it increases dopamine efflux. However, in phase II trials, tesofensine was discontinued for lack of potent effects (Bara-Jimenez et al., 2004; Hauser et al., 2007; Rascol et al., 2008). Unexpectedly, both patients showed significant dose-dependent loss in body weight (Astrup et al., 2008b). Originally, tesofensine was reported to have a greater affinity for dopamine than noradrenaline and serotonin transporters (van de Giessen et al., 2012). However, new studies showed that tesofensine (NS2330) has a lower dopaminergic activity and a stronger effect on norepinephrine: NE (IC50 1.7 nM) > DA (6.5) > 5-HT(11), likewise for the major metabolite M1 (NS2360) NE (0.6) > 5-HT(2.0 nM) > DA(3.0) (Appel et al., 2014; Lehr et al., 2008). Accordingly, we found that tesofensine at therapeutic doses did not induce head weaving stereotypy, a classic behavior induced by most, if not all, dopaminergic-acting appetite suppressants (Kalyanasundar et al., 2020, 2015). Here, we characterized this behavior in more detail for the first time using a subjective visual human observation of a bottom-view videotape and an automatic DeepLabCut analysis. Our human visual inspection revealed that rats performed tongue movements in the air rather than head weaving, which appears to have no apparent function. Hence, it looks like a head weaving stereotypy from a top view, but the tongue protrusions could be seen from a bottom view videotape. This finding suggests that head weaving in rats may be an indirect indicator of tardive dyskinesia-like behavior, which is characterized by oro-facio-buccal-lingual stereotypic movements (Vijayakumar and Jankovic, 2016). Our DeepLabCut analysis found that forward locomotion was reduced in rats that engaged in head weaving stereotypy for hours, as this stereotypic behavior commonly occurred when rats were standing still on all four limbs (**Figures 7A-C**). Our data showed that, unlike phentermine, tesofensine at 2 mg/kg induced few, if any, head weaving stereotypy (**Figure 7C**). However, it reduced forward locomotion (**Figure 7A**), suggesting it slightly affected the overall motor map profile relative to control rats that received saline (see **Figure 7E**, see blue dots slightly separated from control black dots). Our findings of a reduction in locomotion are consistent with a previous study (van de Giessen et al., 2012), although the authors attributed their results to a technical issue. However, we replicated their findings using a more precise method, suggesting that the reduction in locomotion is a genuine phenomenon. Rats treated with tesofensine 2mg/kg commonly transitioned from a sleep-like state to a quiet-awake state. In rare instances, the rats also performed jaw and tongue movements, but at a lower intensity and for brief periods (see **Video 6**).

In contrast, only the higher dose of 6 mg/kg induced strong tongue movements in the air, and this stereotypy exhibited some similarities with phentermine. However, we note it was similar but not identical (**Figure 7E**, green vs. yellow dots). This is expected since tesofensine increases striatal DAT occupancy dose-dependently between 18% and 77% in humans (Appel et al., 2014). Our results suggest that tesofensine at therapeutic doses does not exhibit strong dopamine activity, as evidenced by the absence of head weaving stereotypies. These findings are also consistent with the low risk of abuse for tesofensine, as it has been reported to be unlikely to be abused recreationally (Schoedel et al., 2010).

#### Tesofensine vs. a serotoninergic precursor: 5-HTP

We previously found that the triple combination of phentermine (a dopaminergic compound) plus 5-HTP/carbidopa, a serotoninergic precursor, leads to greater weight loss than each drug alone (Perez et al., 2019). Building on this finding, we investigated whether tesofensine could synergistically interact with 5-HTP/CB to prevent weight loss tolerance. We measured body weight and food intake in the homecages of rats while tesofensine alone or in combination with 5-HTP/CB was administered. Two doses of tesofensine (1 and 2 mg/kg) were used. During the first 6 treatment days, our results demonstrated that this combination did not induce more body weight loss than 5-HTP alone; thus, no synergy effect was observed. However, the group treated with 5-HTP/CB exhibited a marked weight loss tolerance in subsequent days and began regaining body weight afterward. Unexpectedly, tesofensine (at both doses) prolonged the weight loss induced by 5-HTP and completely blocked the body weight tolerance (or body weight rebound) over the following days (**Figure 8A**, see days 7-15). This suggests that tesofensine may have two components: one anorexigenic and a second that stimulates energy expenditure, the latter perhaps mediated by its NE component (Axel et al., 2010; Hansen et al., 2010). This is a promising finding suggesting that tesofensine could be an important adjunct for serotoninergic-acting appetite suppressants to prevent the occurrence of body weight tolerance (Hansen et al., 2010; van de Giessen et al., 2012).

In this regard, a human study found that subjects who took tesofensine for 24 weeks and then stopped taking it for 12 weeks did not regain all their lost weight (Gilbert et al., 2012). Our results support this finding and extend it by showing that tesofensine can also prevent weight rebound after losing weight with another appetite suppressant.

We next tested the interaction of tesofensine and 5-HTP using a variation of the isobolographic assay, with a 1-hour suppression of sucrose intake as the behavioral response. We observed a tendency for a synergistic effect, as indicated by interaction indices lower than 1 of 0.52 and 0.64 for the 1:1 and 3:1 dose ratio, respectively. However, these differences did not reach statistical significance. Therefore, we concluded that tesofensine and 5-HTP/CB exhibited an additive pharmacological interaction for liquid sucrose intake suppression (**Figure 9**). Unlike the synergistic interaction of phentermine/5-HTP/CB (Perez et al., 2019), our results again indirectly suggest that tesofensine, at therapeutic doses, has lower dopaminergic activity. This could explain why we did not find a synergistic effect between tesofensine and 5-HTP.

These experiments also revealed that rats recovered sucrose intake the following day after receiving 5-HTP or tesofensine (**Figure 10**). This suggests that taste aversion does not explain the appetite-suppressing effect of these two drugs. Therefore, tesofensine appears to have anorexigenic properties on its own that are not solely dependent on taste aversion.

### Tesofensine and sweet perception

A human study found that tesofensine increased satiety and decreased cravings for sweet foods after 12 weeks of treatment (Gilbert et al., 2012). However, these effects disappeared after 24 weeks. The reasons for this are unclear (Gilbert et al., 2012). To investigate this further, we used a psychophysical sucrose detection task in rats to determine whether tesofensine affects taste perception. Our data showed that tesofensine did not directly impair the perception of sweetness or its palatability responses (**Figures 10-11**). Although taste responses in rats and humans differ depending on the sweet stimulus, because the homology between sweet receptors in the two species is less than 70% (Nelson et al., 2001), our results suggest that the diminished sweet craving effect observed in humans treated with tesofensine is most likely not due to taste aversion or impairment in sweet perception. Instead, it is likely because of other taste-independent factors, such as post-oral "appetition" signals that mediate food preference via gut-brain nutrient signaling mechanisms (Sclafani, 2013). Further studies are needed to clarify this point.

### Limitations and future directions

The balance of neurotransmitters in the brain, specifically norepinephrine (NE), dopamine (DA), and serotonin (5-HT), is a major determinant of the overall weight loss properties of most appetite suppressants (Baumann et al., 2000; Perez et al., 2019; Rothman and Baumann, 2006). A caveat of our study is that we did not measure the release of these neurotransmitters. Therefore, future studies are warranted to measure NE, DA, and 5-HT simultaneously and map the neurochemical landscape evoked by tesofensine (and other appetite suppressants) using either GRAB sensors with fiber photometry (Feng et al., 2023; Sun et al., 2020) or classic *in vivo* microdialysis with capillary electrophoresis. These studies will clarify the neurochemical profile of each appetite suppressant and will guide us in classifying and combining them better.

## CONCLUSION

Our findings suggest that tesofensine is a promising new therapeutic agent for treating obesity. It modulates neuronal activity in the LH to reduce food intake and promote weight loss. Our data also pave the way for LH GABAergic neurons to be a potential pharmacological target for developing new appetite suppressants to treat obesity. Additionally, this study found that tesofensine may be a valuable adjunct to serotonergic agents to treat obesity, primarily to prevent body weight rebound.

## METHODS

### Drugs and Delivery methods

Several drugs used in the study, including 5-HTP (5-Hydroxytryptophan), carbidopa (CB), and tesofensine, were kindly donated by Productos Medix (Mexico). 5-HTP was dissolved in physiological saline (9% NaCl) at 40°C and injected intraperitoneally. CB was suspended in carboxymethyl cellulose (0.5%) and saline (1:4 ratio) and injected intraperitoneally. Tesofensine was dissolved in saline and injected subcutaneously. Clozapine-N-Oxide (CNO) was dissolved in DMSO/saline (0.1/1ml) and administered intraperitoneally.

### Diets

The control diet of standard chow is composed of 3.43 kcal/g of the following 4.5% lipids, 20% proteins, and 6% carbohydrates (PicoLab, 5053). The High Fat Diet is composed of 4.7 kcal/g of the following: 45% lipids, 20% proteins, and 35% carbohydrates (Research Diet, D12451). Body weight and food intake were measured weekly.

### Subjects: Mice

Male and female adult VGAT-ChR2-EYFP mice (RRID: IMSR_JAX:014548, weighing 20-30 g; n=4) and weaned Vgat-IRES-cre mice (RRID:IMSR_JAX:016962, after 21 postnatal day; n=27) were studied. Both mouse types were purchased from The Jackson Laboratory (Sacramento, CA, USA). VGAT-ChR2-EYFP mice were housed in individual standard laboratory cages with *ad libitum* access to water and chow diet (PicoLab Rodent Diet 20, St. Louis, MO, USA). Vgat-IRES-cre mice were placed in a body weight monitoring protocol (described below). Mice were maintained in standard laboratory conditions of a temperature-controlled (22 ± 1°C) room with a 12:12 h light-dark cycle (7:00 – 19:00). The Institutional Animal Care and Use Committee (CINVESTAV) approved all procedures.

#### Body weight monitoring protocol

Weaned female or male Vgat-IRES-cre mice were separated into groups of 3-5 mice in standard laboratory cages. They were given in their homecages *ad libitum* access to water and either a standard chow diet (PicoLab Rodent Diet 20, St. Louis, MO, USA) or high fat diet (HFD, Research Diet, D12451). Body weights were measured in the evenings once per week for 3 months.

#### Viral constructs

The Cre-inducible adeno-associated virus (AAVs) were purchased from addgene Watertown, MA USA). For ChR2-eYFP mice (AAV5-EfIa-DIO-hChR2(E123T/T159C)-EYFP, #35509, at a titer of 1×1013 vector genome/ml (vg/ml), eYFP-vector (AAV5-EfIa-DIO EYFP), #27056, at a titer of 1.0×1013 vg/ml, and hM4D(Gi) (AAV8-hSyn-DIO-hM4D(Gi)-mCherry), #44362, at a titer of 1.0×1013 vg/ml. Viruses were divided into aliquots and stored at −80 °C before their use.

#### Stereotaxic surgery in mice

After 8-10 weeks of the body weight monitoring protocol, Vgat-IRES-cre mice were individually anesthetized with an intraperitoneal injection of ketamine/xylazine (100/8 mg/kg) and the mice were then placed in a stereotaxic apparatus. A midline sagittal scalp incision was made to expose the skull and two holding screws were inserted into the skull. A microinjection needle (30-gauge) was connected to a 10 µl Hamilton syringe and filled with adeno-associated virus (AAV). Mice were microinjected with AAV (0.5 µl) at a 0.2 µl/min rate. The injector was left in position for 5 additional minutes to allow complete diffusion. For the expressing ChR2, eYFP, or hM4D(Gi). For expressing ChR2, eYFP, or hM4D(Gi) the microinjection was performed unilaterally in the LH (from bregma (mm): AP -1.3, ML ±1.0, and from dura (mm): DV-5.5). In the case of hM4D(Gi), the mice’s head was sutured, and one month was allowed for recovery and expression.

#### Optogenetic experiments

After AAVs infection and to deliver blue light stimulation, a zirconia ferrule of 1.25 mm diameter with multimode optical fiber (200 μm, Thorlabs) was implanted in the LH (from bregma (mm): AP -1.3, ML ±1.0, and from dura (mm): DV -5.3). Mice were allowed one month to recover and to obtain a stable expression of ChR2 or eYFP.

#### Optrode LH implantation in mice

The head of the mice was sutured, and they were allowed 3 weeks for recovery and protein expression. Then, one optrode (16 tungsten wires surrounding a central optic fiber) was unilaterally implanted in the LH (from bregma (mm): AP -1.3, ML ±1.0, and from dura: DV -5.5), and a subcutaneous catheter was inserted subcutaneously for tesofensine administration. Mice were allowed one additional week for recovery.

#### Optogenetic stimulation protocol

A 473-nm laser was intensity-modulated by a DPSS system (OEM laser, UT, USA). Laser pulses were synchronized with behavioral events using Med Associates Inc. software and a TTL signal generator (Med Associates Inc., VT, USA). The patch cord’s optical power was 15 mW and measured with an optical power meter (PM20A, Thorlabs, NJ, USA). However, depending on the efficiency of each fiber, we delivered between 10 and 12.6 mW at the fiber optic tip. Unless otherwise mentioned, the laser was turned on for 2 seconds (at 50 Hz) and off for 4 seconds, with a 10-ms pulse width and a duty cycle of 50%.

#### Open-loop task

Mice were injected subcutaneously with tesofensine (2 mg/kg, 6 mg/kg, or saline) and maintained in their homecage for 30 minutes. After that, mice were placed in an operant chamber (Med Associates Inc., VT, USA) with access to 10% sucrose via licking a central sipper port. Mice received a 5-min block of no stimulation (off) followed by the delivery of the optostimulation pattern (on) for 30 minutes (see *Optogenetic stimulation protocol* section). All licking responses were recorded by a contact lickometer (Med Associates Inc., VT, USA).

Chemogenetic silencing

Five mice fed a high-fat diet (HFD) with the expression of hM4D(Gi) were placed in their homecage with an automated feeder (FED 3, Open Ephys; Lisbon, Portugal). Each time mice executed a nose poke, the feeder delivered a chocolate pellet (20 mg, #F05301, Bio-Serv, Nueva Jersey, USA). Tesofensine (2 mg/kg, subcutaneously), Clozapine-N-Oxide (CNO, 3 mg/kg, intraperitoneally), both drugs or vehicle (DMSO or saline) were administered at 18:30 (30 minutes before the dark cycle begins). The vehicle was always administered in the subsequent session to avoid a carryover effect of drugs. Sessions were counterbalanced between subjects.

#### Electrophysiology in mice

##### Extracellular recordings of LH in freely moving Vgat-IRES-cre mice

Multi-unit recordings were performed using a Multichannel Acquisition Processor system (Plexon, Dallas, Texas) and a Med Associates (Fairfax, VT) interphase to record behavioral events simultaneously. Voltage signals were sampled at 40 kHz and digitalized at 12 bits resolution. Action potentials with a signal-to-noise ratio larger than 3:1 were analyzed. They were identified online using a voltage-time threshold window and a three-principal component contour template algorithm (Gutierrez et al., 2010). After the experiment, spikes were sorted using Off-line Sorter software (Plexon Inc.).

Mice were recorded in a baseline period of 5 min, then saline (0.15 ml, 0.9 %) or tesofensine (2 mg/kg) was administered via a subcutaneous catheter. Thirty minutes after, mice received optogenetic stimulation in blocks of 5 min Off-5 min followed by a block On (2s on-4s off) for 40 min (eight blocks total).

##### Histology in mice

Mice were anesthetized with sodium pentobarbital (75 mg/kg), then perfused intracardially with PBS 1x and paraformaldehyde at 4%. Their brains were removed and stored in 4% paraformaldehyde solution for 48-h hours and put in a 30% sucrose solution for 72-h hours. The brain was sliced, and sections of 40 μm were mounted in Dako fluorescence mounting medium. Immunofluorescence was observed using a ZEISS LSM 800 confocal microscope.

### Subjects: Rats

Adult male Wistar rats were used in all experiments: From weaning thirty-three rats were housed in pairs until 12 weeks of age, after which they were housed individually. During those 12 weeks, the rats were given either a high-fat diet (HFD) or a chow diet and had *a*d libitum access to water. Six male rats (n=3 with HFD, n=3 with chow food) had electrodes implanted in their LH for electrophysiological recordings.

A total of 24 male rats, 330–370 g, were used for the locomotion and stereotypy experiments. The rats were housed in their home cages in pairs and had *ad libitum* access to food and water. In the acrylic box, rats were given one pellet of chow and one high-fat diet (HFD) attached to a side wall. In the isobologram studies, lean rats (n=142) were maintained on a chow diet and received 10% sucrose instead of water for one hour during the treatment. Finally, 4 male rats were trained in a sucrose detection task (data not shown). Unless otherwise mentioned, all rats were kept in a temperature-controlled environment at 21 ± 1°C and on a 12:12 hour light-dark cycle (06:00-18:00).

#### Tesofensine administration on a diet-induced obesity model

This study investigated the effectiveness of tesofensine in treating obesity using a diet-induced obesity model (Gajda, 2008; Kimura et al., 2017). Three-week-old, weight-matched male rats (60-65 g) were housed in pairs and fed either a standard chow diet or a high-fat diet until they reached 12 weeks of age. After 12 weeks, the rats were divided into four groups: Chow-Saline (n=6), Chow-Tesofensine (n=7), HFD-Saline (n=6), and HFD-Tesofensine (n=8). Rats were housed individually during the treatment and maintained access to their assigned diet. The two tesofensine groups received 2 mg/kg tesofensine, injected subcutaneously 60 minutes before the dark cycle onset for 15 consecutive days. The rat’s body weight and food intake were measured every 24 hours and expressed as daily body weight gain compared to the first day of drug administration. After the treatment, the rats were fasted for 12 hours and euthanized by an overdose of pentobarbital sodium anesthesia (100 mg/kg i.p.). Total fat mass was measured by weighing the gonadal, perirenal, and mesenteric adipose tissues (Kim et al., 2000; Schipper et al., 2018; Wallenius et al., 2002).

#### Surgery for extracellular recording in rat’s LH

The electrophysiological data was collected and processed as detailed in extracellular recordings in mice. All rats underwent surgery under anesthesia, obtained by an intraperitoneal injection of xylazine (8 mg/kg) and ketamine (80 mg/kg). A local analgesic, lidocaine (4 mg/kg of 1% solution), was administered subcutaneously under the head skin. The rats were then placed in a stereotaxic apparatus for implantation of a homemade electrode array composed of 16 tungsten wires (35 μm in diameter, arranged in a 4×4 array with an area of 1 mm2) in the right LH (AP -2.1 mm, ML -1.5 mm from bregma, and DV -8.3 mm from the dura). The electrode array was attached to a dedicated tungsten filament inserted into the LH, and a stainless-steel screw was soldered to a silver wire for electric ground, which was screwed above the cerebellum and cemented into the skull.

For subcutaneous catheter implantation, the rats underwent two small incisions (∼1mm) in the superior left abdomen and dorsal neck areas. Sterilized silicone tubing (12 cm long, Silastic laboratory tubing, Dow Corning, Midland, MI, CAT. No. 508-004) was used as a catheter and tunneled subcutaneously from the back incision to the dorsal neck incision. The incisions were then sutured and closed using USP 3-0 (Atramat, Mexico). After surgery, the rats were treated with intraperitoneal enrofloxacin (10 mg/kg) and meloxicam (2 mg/kg) for three consecutive days. Experiments began seven days after surgery(Perez et al., 2019).

#### Histology in rats

Rats were anesthetized with an overdose of sodium pentobarbital (150 mg/kg), then perfused intracardially with PBS 1x and paraformaldehyde at 4%. The brain was removed and put it in a 10% sucrose solution for 24 h, followed by sequential increases in sucrose concentration until reaching 30% in a 72-h period. The brain was then sliced coronally (50 μm thick) and stained with cresyl violet. For histological confirmation of electrode location in the brain, the electrodes were covered with DiI lipophilic carbocyanine dye (1%; Sigma-Aldrich) allowing the observation of the fluorescent track left by the electrodes.

#### Chronic treatment with tesofensine and 5-HTP/CB

The interaction between tesofensine and 5-HTP was characterized in lean male rats fed a standard chow diet. The rats were given daily injections of tesofensine plus a fixed dose of 5-HTP (31 mg/kg, i.p.) and CB (75 mg/kg, i.p.) for 15 days, following one of six protocols: vehicle (control, n=6 for each group), 5-HTP/CB (31 and 75 mg/kg), tesofensine1 (1 mg/kg, s.c.), tesofensine2 (2 mg/kg, s.c.), tesofensine1 + 5-HTP/CB, and tesofensine2 + 5-HTP/CB. CB was administered 30 minutes before 5-HTP, and tesofensine was injected subcutaneously 30 minutes after 5-HTP. Body weight and food intake were measured every 24 hours and expressed as daily body weight gain or food intake relative to the first day of drug administration. The drugs were administered between 13:00 and 18:00 h.

#### Isobologram curve for sucrose intake assay

In order to investigate the potential pharmacological interaction between tesofensine and 5-HTP/CB, we developed a novel isobolographic assay based on 10% sucrose consumption (Tallarida, 2000). Before the experiment, rats were only fed a chow diet and trained to lick a bottle filled with 10% sucrose for one hour each day for six days, during which they were habituated to regular sucrose consumption. After establishing baseline sucrose intake levels, the rats were randomly divided into groups of n=6, matched for their initial sucrose intake levels (basal period). On the seventh day (experimental day), the rats received either a vehicle or a single dose of tesofensine (0.1-5 mg/kg, s.c.), 5-HTP (0.5-100 mg/kg, i.p.), or a combination of tesofensine and 5-HTP at a 1:1 or 3:1 ED30 ratio. For tesofensine alone, the sucrose bottle was presented one hour after administration. For 5-HTP alone, the group received a vehicle or CB (75 mg/kg i.p.) 30 minutes before 5-HTP, and the sucrose bottle was presented 90 minutes after the last injection. CB was given first for the combination drug curve, followed by 5-HTP, and tesofensine was administered 30 minutes later. The sucrose bottle was always presented one hour after the last injection. The total sucrose intake was measured to the nearest milliliter, and the sucrose bottle was available for one hour between 17:00 to 18:00.

#### Homegustometer appararatus

A homegustometer is a homemade behavioral setup that adapts the animal homecage for psychophysical tasks. Rats can perform the task day and night for weeks. To create the homegustometer, the frontal wall of a standard homecage was opened to allow the insertion of three sippers. The central sipper was fixed in position, while the two lateral sippers were adjusted to 60 degrees relative to the central sipper. A metal floor was inserted into the homecage’s floor and served as ground for the contact lickometers. All licks were recorded using a Med Associates interphase (Fonseca et al., 2018).

#### Sucrose Detection Task

To assess sucrose’s perception, rats were trained to visit a central port and give between 2 and 5 licks in an empty sipper to receive a 10 µL drop comprising either water or one of five sucrose solutions with varying concentrations (0.5, 1.3, 3.2, 7.9, or 20% w/v). Trials were balanced such that the probability of receiving water (0%) or sucrose (any concentration) was 0.5, and they were presented in pseudo-random order. Then the subjects were required to report whether the drop contained or did not contain sucrose, by approaching and then licking the left outcome port if the stimulus was water (0%), and the right port if it was sucrose. This rule was counterbalanced across subjects (Fonseca et al., 2018). Successful detection led to reward, which consisted of the delivery of a drop of water per each of the subsequent three licks. Error trials were unrewarded. Trials ended 0.3 seconds after the last water drop for rewarded trials; and for unrewarded trials, the trials ended 0.3 seconds after the first dry lick. After receiving either the Stimulus or the Reward, the subjects could keep dry licking the ports with no penalties but wasting time to complete more trials and obtain more rewards. The number of dry licks after the Stimulus in the central port is an indirect measurement of the hedonic value of the tastant; indeed, in our task the post-stimulus licks increased with sucrose palatability(see Perez et al., 2013). For this reason, the task could measure oromotor palatability responses elicited by one single drop of sucrose.

The subjects were trained in the homegustometer which allowed a 23 h training in their homecages. Four rats were trained in the sucrose detection task for at least 10 days until they achieved 75% correct responses during two consecutive days. Following initial training (not shown), rats underwent 1-2 months break before being reintroduced to the homegustometer. Three baseline sessions were recorded before 5 days of treatment with s.c. tesofensine 2 mg/kg. This was followed by 3 days of saline injections in the post-tesofensine period. The equipment was cleaned and recalibrated daily at noon, and injections occurred around 18:00. Thus, rats had access to water *ad libitum* in their home cages for one hour while the homegustometer was being cleaned.

#### Open acrylic box and locomotor activity

For behavioral experiments, locomotor activity was measured in an acrylic box (41.5 cm in length, 30 cm in width, and 26 cm in height) coupled with a camera (in the bottom view position). From a bottom-view video recording, the animals’ position at x and y coordinates of rats’ noses, forelimbs, hind-limbs, and tail base was tracked using DeepLabCut software (DLC)(Mathis et al., 2018). A video was recorded at 60 frames per second (fps) with a resolution of 1280 x 720 pixels using a Kayeton camera (model KYT-U400-MCS2812R01). The forward locomotion was tracked using the rats’ center mass of the hind-limbs method and plotted as total distance traveled (cm) for 240 minutes.

### Data analysis

All data analysis was performed using MATLAB (The MathWorks Inc., Natick, MA), GraphPad Prism (La Jolla, CA, USA), DeepLabCut, and Python. For isobologram analysis we wrote a custom Matlab script that is available as supplementary material. Unless otherwise indicated, we used the mean ± sem and the α level at 0.05.

#### DeepLabCut

DeepLabCut 2.2.3 was used to track all points of interest in a bottom-view videotape. A network was trained using 15 frames from 12 randomly selected videos for 1,030,000 iterations. The learning rate was decreased in a stepwise fashion, starting at 0.005 for 10,000 iterations, then 0.02 for 750,000 iterations, 0.002 for 800,000 iterations, and finally 0.001 for the remaining 103,000 iterations. Ten outliers from each training video were then corrected by relabeling points with a likelihood below 0.9. The network was then refined using the same number of iterations.

#### Measurements of head weaving stereotypy

The head weaving stereotypy was measured using the data obtained from DLC tracking of the angular variation of the Euclidean position of the nose regarding its base tail. Snippets were made from the angular variation data by averaging 3600 data points corresponding to one minute of the session time. Outliers were removed, and the first derivative of the snippet data was taken. Once this new vector was obtained, the standard deviation of the data was calculated, and it was found that the values greater than two negative standard deviations and less than two standard deviations and the centroid corresponding to the hind-limbs remain static (variation less than 1cm between minutes) corresponds to data describing the head weaving behavior. We consider stereotypy only for moments in which the rat remained immobile with four legs in contact with the floor (Perez et al., 2019). These results were displayed as the percentage of time spent in each behavioral state.

#### Isobologram analysis

To evaluate the pharmacological interaction between tesofensine and 5-HTP, an isobolographic analysis was performed using a sucrose intake assay. The data were presented as the mean ± standard error of the mean (SEM), with six rats per group. Dose-response curves were plotted by showing the percentage of anorexigenic effect against the log dose. The percentage of the anorexigenic effect was calculated by subtracting the basal intake and then dividing the volume of sucrose consumed in one hour by the following formula:

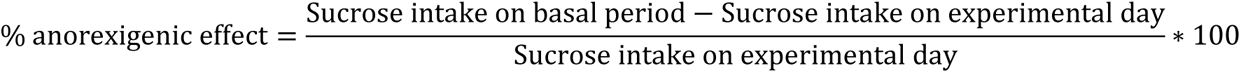

#### Statistical analysis

For statistical analysis of differences between groups treated with tesofensine plus 5-HTP/CB, a repeated measure two-way ANOVA was conducted, followed by a Tukey’s *post hoc* test. The experimental Effective Dose (ED30) for tesofensine and ED30 for 5-HTP were found using linear regression of the log dose-response curves for the isobolographic analysis. The theoretical additive line was constructed by plotting the ED30 values of tesofensine and 5-HTP alone and by computing the theoretical ED30 of the combination. The dose-response curve was obtained after the co-administration of fixed doses of tesofensine and 5-HTP based on fractions (1/2, 1/4, and 1/8) relative to their respective ED30 values. A one-way ANOVA followed by a Dunnett’s test post-hoc was used in the GraphPad Prism 6 software.

The dose ratios of tesofensine and 5-HTP were 1:1 and 3:1 and the experimental ED30 values of the combination were determined from the log dose-response curve for each dose ratio using linear regression. Statistical difference between each theoretical ED30 and its experimental ED30, along with confidence intervals for each dose ratio and its interaction indices, were calculated following the modified t-test proposed by (Tallarida, 2000), implemented in a homemade MATLAB script (available as supplementary material).

#### Electrophysiological data analysis

Statistical analysis and figures were performed using MATLAB (MathWorks). We used a chi-square test to assess differences in the proportion of neurons recruited. The Peri-stimulus Time Histograms (PSTHs) were then color-coded into z-scores.

t-distributed Stochastic Neighbor Embedding (t-SNE) is an automatic dimensionality reduction method that tries to cluster similar firing rates in a low-dimensional space to optimally preserve neighborhood identity (Maaten and Hinton, 2008). In this manuscript, we used t-SNE to cluster neurons with similar firing rate modulation into neuronal ensembles (Coss et al., 2022). We also used t-SNE to analyze the profile of motor effects induced by appetite suppressants, in this case, clustering rats exhibiting similar motor side effects.

#### CRediT authorship contribution statement

##### CIP

Conceptualization, Data curation, Software, Formal analysis, Supervision, Validation, Investigation, Visualization, Methodology, Writing – original draft, Writing – review & editing. **JLI**, Conceptualization, Data curation, Software, Formal analysis, Investigation, Writing – original draft. **AL**, Data curation, Software, Formal analysis, Investigation, Writing – original draft, **XD** Methodology, Investigation, **OM** Methodology, Investigation, **BA** Methodology, Software, Formal analysis, **MGM** Investigation, **EGL** Visualization, Data curation, **EF** Conceptualization, Methodology,

Software, **GC** Software, Supervision, **RG** Conceptualization, Resources, Software, Supervision, Funding acquisition, Validation, Visualization, Methodology, Writing – original draft, Project administration, Writing – review & editing.

## Acknowledgments

We thank Professor Sid A. Simon for insightful comments on this manuscript. We also thank Ricardo Gaxiola, Victor Manuel García Gomez, and Fabiola Hernández Olvera for their invaluable animal care.

## Supplementary Figures

**Figure 2 supplementary 1.**
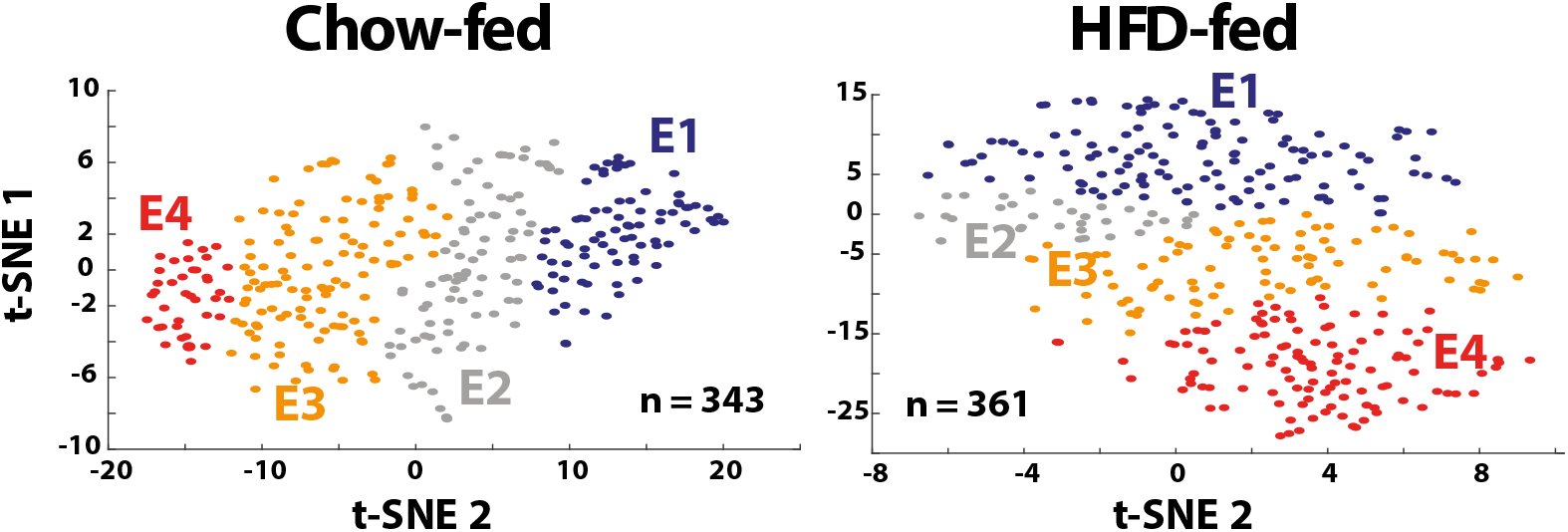
T-SNE analysis uncovered four ensembles responding to tesofensine. t-SNE analysis of firing rates was used to group similar activity patterns into ensembles. Neurons were assigned into four ensembles (E1-4) for Chow-fed rats (left) and HFD-fed rats (right). Each dot represents a single neuron and the color of the ensemble it belongs in the t-SNE map.

**Figure 9 supplementary 1.**
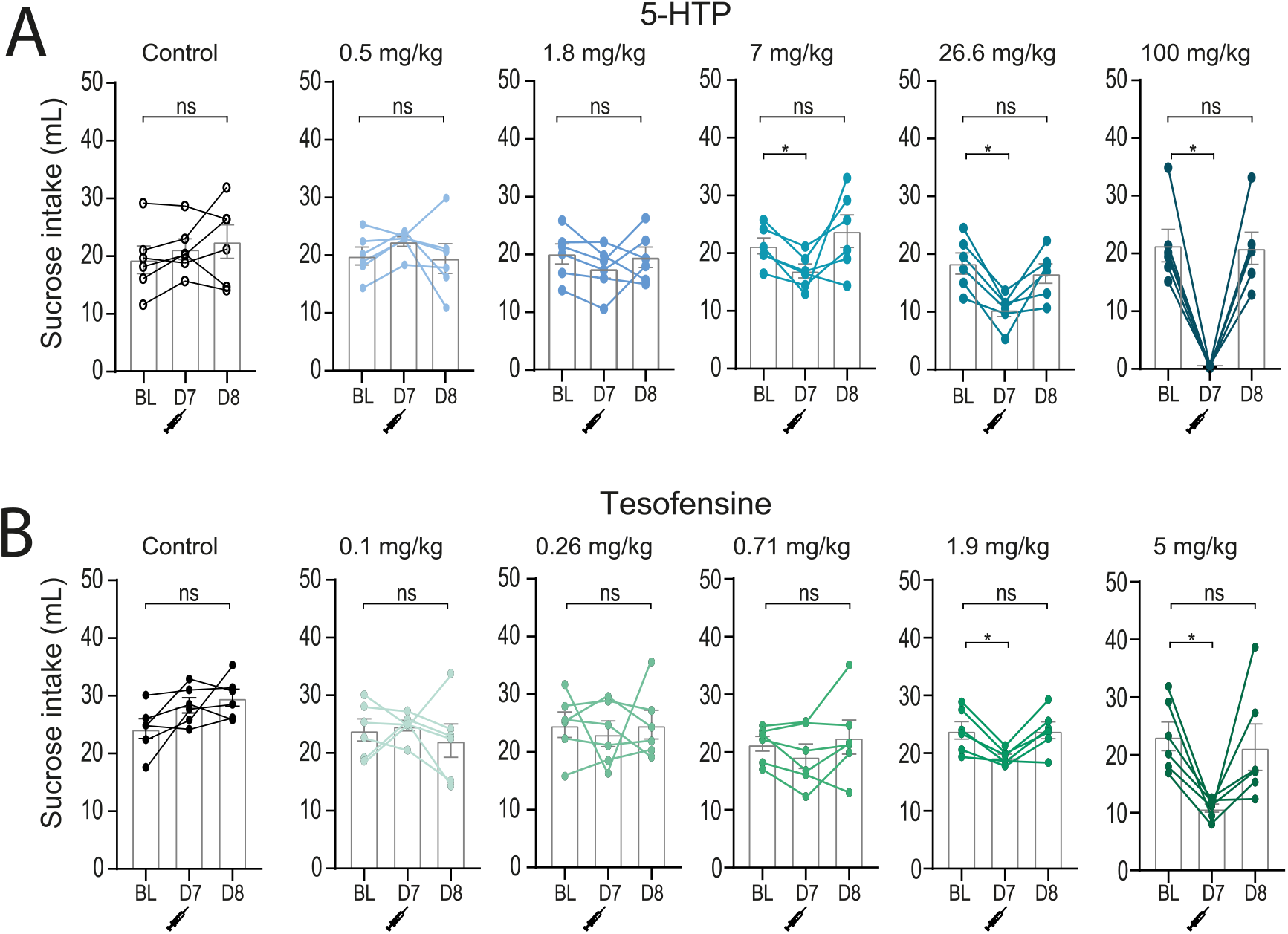
The appetite suppressant effects of tesofensine and 5-HTP/CB is not due to taste aversion. A. The graph plots the average 1-hour sucrose intake during baseline sessions. On day 7 (D7), drugs were administered before giving access to sucrose. A dose-dependent intake suppression can be seen. Finally, on day 8 (D8), intake was observed the day after treatment. 5-HTP/CB suppressed acute sucrose intake on day 7 (D7), but consumption returned to baseline levels on day 8 (D8). Bar plots show acute sucrose intake (1 hour) for control rats (vehicle injection) and treated rats (different doses of 5-HTP/CB). Each dose was tested only once in a new set of naïve rats. Note that 5-HTP/CB induced a dose-dependent decrease in sucrose intake. BL = baseline sucrose intake. **B.** Same convention as panel “**A**” but for tesofensine. * p<0.05 significantly different.

